# Genome-wide Pervasiveness and Localized Variation of *k*-mer-based Genomic Signatures in Eukaryotes

**DOI:** 10.1101/2025.06.12.659337

**Authors:** Niousha Sadjadi, Camila P. E. de Souza, Gurjit S. Randhawa, Kathleen A. Hill, Lila Kari

**Author notes:** Joint senior author.

## Abstract

Genomic signatures are taxon-specific patterns in nucleotide sequence composition observed across different regions of a genome, used in the taxonomic classification of organisms and in inferring their evolutionary relationships. However, the nature and extent of the pervasiveness of a genomic signature across the expanse of a Telomere-to-Telomere (T2T) assembly, especially across the functionally diverse sequence elements and highly repetitive regions, remain counterintuitive and underexplored. This study aims to bridge this knowledge gap by systematically investigating the pervasiveness and variation of the genomic signature across the human genome and the genome of each of three other eukaryotic species from different kingdoms. Using the alignment-free *k*-mer-based *Frequency Chaos Game Representation (FCGR) of DNA sequences*, this study qualitatively and quantitatively analyzes the variations of the genomic signature along an entire genome. Qualitative analysis is first performed through visual inspection of FCGR patterns across different chromosomes of a species. In parallel, a quantitative analysis evaluates the variation of the genomic signature within a genome by comparing eight distance measures to identify the optimal one for the datasets in this study. By taking an intragenomic perspective with detailed analysis of chromosome landscapes these analyses reveal that, while the genomic signature is preserved in most genomic regions, exceptions exist in localized regions, such as tandem repetitions of short and long repeat units. Upon determining this pervasiveness, we assemble novel pipelines aimed at selecting a short contiguous representative genomic segment that encapsulates the sequence composition patterns characteristic of the entire genome. These representative segments are then used to assess intragenomic variation of the genomic signature, demonstrating that only a small proportion of segments (namely those characterized by regional density of short and long tandem repeats) show high distance values from the representative. Notably, in the human genome, 80% of the segments have a distance of less than 0.24 (on a [0,1] DSSIM scale) from the representative. Moreover, we demonstrate that using these representative segments improves down-stream tasks, e.g., increasing one-nearest-neighbor (1-NN) taxonomic classification accuracy by 7% compared to selecting a random genomic segment to serve as a proxy of the genome. Lastly, this study presents a special-purpose graphical user interface (GUI) software tool, *CGR-Diff*, designed to provide both visual and quantitative comparisons of FCGRs of sample or user-provided DNA sequences, thereby facilitating intragenomic variation analysis of genomic signature within and across species.

## Introduction

Recent advances in long-read sequencing and sequence assembly algorithms have led to many Telomere-to-Telomere (T2T) genome sequence assemblies for various species [38, 56], including human, uncovering significant heterogeneity and structural variation within the genome [56]. While it is known that across the genome there are several structural elements (e.g., telomeres, centromeres, and chromosome arms) which differ in function and composition [64], analysis of T2T sequences has revealed additional insights. Beyond the structural variation that exists within the genome, several factors vary along the length of chromosomes, including GC content, gene density, and the distribution of long interspersed repeat elements (LINEs), long terminal repeats (LTRs), microsatellites, and other repeats [38, 64, 81]. Furthermore, some studies have shown that even regions traditionally considered highly uniform in sequence composition (e.g., Internal Transcribed Spacers (ITS) and rDNA) exhibit extensive variation across different parts of the genome [13, 75].

In spite of this regional diversity in genome sequence composition associated with local functional attributes, some studies have suggested the presence of certain global characteristics (e.g., dinucleotide relative abundance [11, 30, 35], or the relative frequency of oligonucleotides [18]) that are repeated across different segments of a genome and are referred to as the genomic signature [35]. The studies introducing these concepts hypothesized that the genomic signature is pervasive genome-wide and species-specific, although this conclusion was based on limited analyses of short genomic regions (50 kbp to 100 kbp) [11, 30, 35] or a small number of species [18]. In spite of the small scale of these studies, the assumption of the pervasiveness of genomic signatures within a genome took hold and has been successfully used in numerous applications, including phylogenetics and evolutionary analyses [17, 46], taxonomic classification [18, 33, 34], genome adaptation studies [6], the identification of emergent pathogens [29], and the classification of cancer genomes [47].

Prior work has identified various quantitative approaches that could serve as a genomic signature [9, 53, 76]. One of the earliest approaches was the use of Dinucleotide Relative Abundance Profiles (DRAPs) [14, 30, 35–37], which calculates the ratio of the observed frequency of a dinucleotide (a pair of consecutive nucleotides) to its expected frequency (i.e., the product of the frequencies of its constituent nucleotides). The DRAP concept was then generalized to the oligonucleotide relative abundance profile of a DNA segment, computed as the ratio of the observed frequency of an oligonucleotide to its expected frequency [76]. The Generalized Genomic Signature [5] is a similar approach that operates by first filtering out the background nucleotide composition and then measuring only the deviation of oligonucleotide frequencies from this background composition.

Another approach to construct a genomic signature is through the Chaos Game Representation (CGR) [28] of DNA sequences and its derivative, Frequency Chaos Game Representation (FCGR) [3, 19]. The CGR of a DNA sequence is a two-dimensional binary image whereby each pixel represents the presence/absence of an oligonucleotide in the sequence [33], and where the resolution of the image determines the oligonucleotide length [53]. Thus, the CGR of a DNA sequence is a simultaneous representation of the distribution of oligonucleotides of a certain length within that sequence. FCGR generalizes CGR by providing a quantitative view: In an FCGR of resolution 2^*k*^ ×2^*k*^, where *k* corresponds to the length of the oligonucleotide [76] (also referred to as a *k*-mer), the intensity of each pixel represents the frequency of a specific *k*-mer within the sequence, making the entire plot a comprehensive visual representation of *k*-mer frequencies in the originating DNA sequence. FCGRs of DNA sequences exhibit fractal geometric patterns, and the genomic signature represented by an FCGR is correlated to other methods such as DRAP, in that DRAP can be deduced from FCGR [53, 76], but not vice versa [76]. CGR and FCGR have gained significant attention through their usability in bioinformatics [32, 46], due to their ability to visually encapsulate genome-wide sequence composition patterns, and given their robustness, flexibility, and applicability to sequences of any length [18]. Specifically, FCGR has been effectively used for alignment-free genome comparisons in taxonomic analysis [1, 2, 43], thanks to the computational efficiency that results from its alignment-free nature, which allows bypassing the computationally expensive step of multiple sequence alignment [2].

While the aforementioned studies have suggested that a *k*-mer-based genomic signature is pervasive across the genome of an organism [17, 30, 46] and this assumption has been successfully used in various bioinformatics applications, the extent to which this pervasiveness holds throughout a T2T genome assembly remains counterintuitive [17] and underexplored. In particular, the intragenomic variability of such a genomic signature across different genomic regions still awaits a comprehensive investigation. Should the hypothesis of genomic signature pervasiveness be conclusively proven, an alignment-free genome comparison algorithm would still depend on a method to reliably select a DNA genomic segment that reflects the nucleotide composition characteristics of the whole genome. Given the observed heterogeneity and structural variations within a genome and the expected impact of this heterogeneity on the genomic signature, finding such a representative DNA genomic segment could be challenging.

This study aims to fill these gaps by providing extensive qualitative and quantitative analyses supporting the hypothesis that a *k*-mer-based genomic signature is largely preserved along the length of each individual T2T genome in human and other eukaryotic species, with notable exceptions and localized variations. It also proposes two computational pipelines for selecting a short representative DNA segment that captures the nucleotide composition of the entire genome, at a given resolution (*k*-mer size), and that can be used to explore the sequence composition variability along a T2T assembly. Finally, through several computational experiments, this study demonstrates that short representative genomic segments can be successfully used in downstream tasks such as taxonomic classification.

Concretely, to extract genomic signatures that reliably encapsulate the characteristics of the genome, this study uses FCGR, which allows both visual and quantitative comparisons of genomic signatures. Visual inspection of FCGR images from each chromosome of a T2T genome reveals consistent overall patterns, although variations in image intensity are observed. Beyond a qualitative analysis, this study shows that when comparing FCGRs quantitatively, some distance measures outperform others in capturing notable biological differences in the genomic signature. Through various intra- and intergenomic experiments, the Structural Dissimilarity Index (DSSIM) is suggested as a suitable measure for FCGR comparison for the datasets in this paper.

Using DSSIM, the *Representative Segment Selection Pipeline* (RepSeg) is then introduced as an algorithm to identify a genomic segment whose FCGR has the minimum average distance to those of other segments within the same genome. To increase the computational efficiency of RepSeg, a computationally optimized pipeline, the *Approximate Representative Segment Selection Pipeline* (aRepSeg) is also proposed: Depending on the application, RepSeg can be used when high accuracy is prioritized, while aRepSeg can be used for datasets with large genomes or for time-sensitive applications. The representative segment identified by either pipeline encapsulates genome-wide nucleotide compositions and enables quantitative investigation of intragenomic variation by measuring the distances between constituent genome segments and the representative segment, thereby revealing longitudinal changes in the genomic signature. Also, the representative segment serves as an effective proxy for downstream tasks such as taxonomic classification, and in this study, a benchmark dataset is designed to demonstrate that using representative segments as training samples improves the performance of a one-nearest-neighbor (1-NN) classifier by 7%.

Finally, to facilitate both visual and quantitative comparisons of FCGRs derived from different genomic sequences, and to support the replication of the analysis in this study, a software tool with a graphical user interface (GUI), called *CGR-Diff* is developed. Unlike existing general-purpose GUI tools for intragenomic analysis, such as Integrative Genomics Viewer (IGV) [74], UGENE [57], VDAP-GUI [51], and GEMINI [59], or alignment-free *k*-mer-frequency comparison tools such as KAST [71] and TreeWave [12], *CGR-Diff* is specialized for FCGR-based analysis of the intragenomic variation. The software visualizes the FCGRs of two independent sequences, whether from the same species or different ones, highlights their differences, and provides several quantitative methods to measure their dissimilarity.

The main contributions of this study are:

- Extensive qualitative and quantitative evidence of the genome-wide pervasiveness and localized variation of a *k*-mer-based genomic signature in eukaryotes;
- Design and implementation of two computational pipelines for the selection of a short contiguous representative DNA segment (500 kbp) to act as genome proxy for taxonomic and other analyses, enabling a comprehensive quantitative assessment of the intragenomic variability of the genomic signature. For example, our results show that across the human genome, 80% of genomic segments have a DSSIM distance of less than 0.24 from the representative (on a [0,1] scale);
- Development of *CGR-Diff*, a novel software tool with a graphical user interface (GUI) for the visual and quantitative comparisons of FCGR-based genomic signatures of sample or user-provided DNA sequences.

To the best of our knowledge, this is the first study to examine the pervasiveness of FCGR-based genomic signatures across whole genomes of eukaryotic species from four different kingdoms of life, and propose pipelines for selecting a representative DNA segment that preserves the key characteristics of the whole genome.

## Methods

This section first describes the dataset and genome sequences utilized in this study. Next, it provides an overview of FCGR, a graphical representation of genomic signatures used in this paper. This is followed by a detailed description of various distance measures employed to compare FCGRs, which serve as a key component of the proposed pipeline. Then, RepSeg is introduced as a pipeline designed to select a short-length representative of the entire genome that can act as a proxy of that genome for computational analyses. To further enhance computational efficiency and reduce memory usage, aRepSeg, an optimized version of RepSeg, is proposed to select representative genome segments more efficiently. Subsequently, *CGR-Diff*, a novel graphical software developed in Python, is described as a tool that enables the visualization and comparison of genomic FCGRs and facilitates both intragenomic and intergenomic studies. The software allows users to either upload two genomic FASTA files or select segments from predefined genomic assemblies of various species, as listed in Fig. 1b. After uploading or selecting sequences, users can identify and select specific segments from each sequence for FCGR comparison. Additionally, *CGR-Diff* offers built-in tests for analyzing intersegment variation, increasing its applicability in genomic research. Finally, the experiments conducted in this study are outlined (see Fig. 1 for an overview of the experiments and dataset).

**Figure 1:**
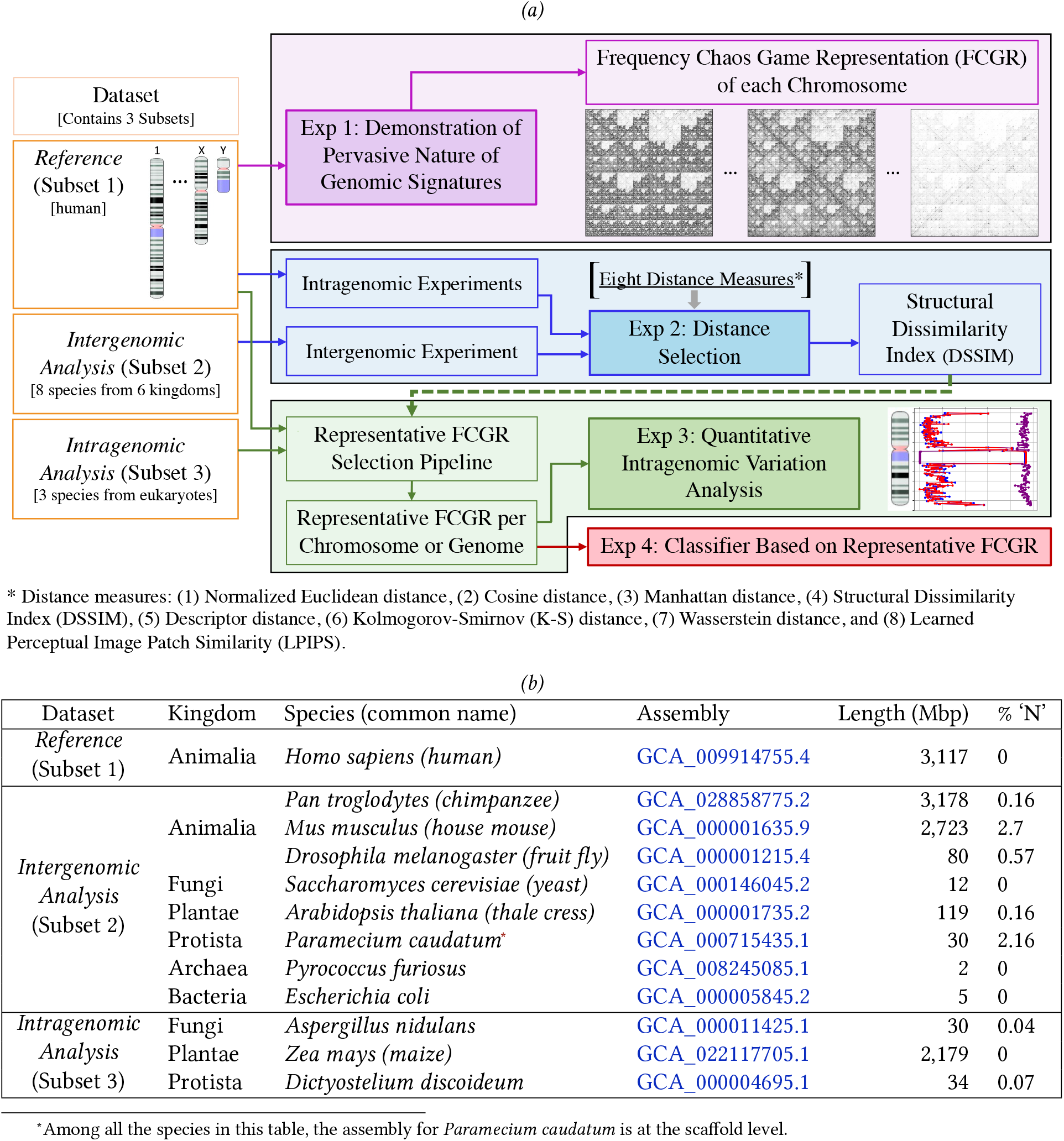
Method overview and dataset. **(a)** Overview of the four experiments and their interrelationships: Experiment 1 is an independent study exploring the pervasiveness of genomic signatures across chromosomes within a single species. Experiment 2 conducts internal tests to identify the most appropriate distance measure for comparing genomic signatures. Experiment 3 applies the selected distance measure from Experiment 2 to analyze intragenomic variation across the entire genome through representative segment selection. Experiment 4 assesses the effectiveness of the representative segments suggested by pipelines using a 1-NN classifier. **(b)** Summary of selected species: This includes a list of the selected species for our subsets, detailing their GenBank assemblies from the NCBI database, genome lengths, and the percentage of unknown nucleotides (represented as ‘N’) in their genomes.

### Dataset

The DNA sequence dataset consists of three subsets covering the whole genomes of several model species from different kingdoms creating a comprehensive resource for various experiments. The human genome, functioning as the *Reference Subset* (Subset 1), serves as the core and primary data in the study. This choice is justified by the structural and functional complexities of the human genome, along with the extensive annotations and supplementary information available for its Telomere-to-Telomere (T2T) assembly [56, 64]. In addition to the human genome, the dataset includes the whole genomes of eleven other species, organized into two different subsets: the *Intergenomic Analysis Subset* (Subset 2), and the *Intragenomic Analysis Subset* (Subset 3). These genomes are specifically selected to be used alongside the human genome for specific parts of the experiments and analyses. The subsequent paragraphs provide details of these genomes, and indicate the source to retrieve their data as well as annotations, in cases where the annotations are available. Additionally, our GitHub page includes the data assembly files for each species, instructions on how to retrieve annotations, and the final annotation files.

To retrieve the human genome, the latest assembly, “T2T-CHM13v2.0,” released in 2022 [56, 64], was downloaded from the NCBI database (link in Fig. 1b) on May 9, 2023. This fully annotated assembly, developed by the Telomere-to-Telomere (T2T) consortium [73], offers a gapless and complete representation of the human genome, covering all chromosomal regions. It contains all known structural variations, including tandem repeats for each chromosome, and features a fully colored and annotated ideogram. The annotations for the different parts of the chromosomes are obtained from GitHub, the NCBI database, or are manually extracted from the NCBI Genome Data Viewer [54] on Dec 4, 2023.

The *Intergenomic Analysis Subset* (Subset 2) includes the genomes of eight species with different phylogenetic or evolutionary distances from the human genome, which are used for the distance selection experiment. Two of these species, *Pan troglodytes* and *Mus musculus*, are selected from Class Mammalia due to their common origin from an ancestral eutherian genome, which results in strong resemblance in chromosome structures and gene sequences [24]. In addition to the mammalian genomes, six other species are selected for Subset 2, each representing one of the six kingdoms of life: *Drosophila melanogaster* (Animalia), *Saccharomyces cerevisiae* (Fungi), *Arabidopsis thaliana* (Plantae), *Paramecium caudatum* (Protista), *Pyrococcus furiosus* (Archaea), and *Escherichia coli* (Bacteria). The genomes of these species differ significantly from the human genome in terms of karyotypes and gene density. Notably, *Pyrococcus furiosus* (Archaea) and *Escherichia coli* (Bacteria) are prokaryotes, meaning their DNA is not encapsulated within a nucleus, and the compactness and regulatory mechanisms of their genomes differ substantially from those in eukaryotes. These eight species are chosen to span a range of genomic resemblance to the human genome, enabling the evaluation of whether these differences are reflected in the FCGR distance analysis. The assemblies for these species were retrieved on May 25, 2024 from the NCBI database (link in Fig. 1b). Among them, the assemblies of *Saccharomyces cerevisiae* (Fungi), *Pyrococcus furiosus* (Archaea), and *Escherichia coli* (Bacteria) are gapless and complete, containing no unknown nucleotides (‘N’), while the rest of the assemblies include ‘N’ within their genome. In generating FCGRs, these unknown nucleotides are removed without introducing *k*-mer artifacts by discarding *k*-mers that contain ‘N.’ Moreover, among these eight species, all assemblies are at the chromosome level, except for *Paramecium caudatum*, whose assembly is at the scaffold level. Fig. 1b includes the additional information of these species including their assembly number in the NCBI database, length of their genome sequence, and the percentage of unknown nucleotides in their assembly.

The *Intragenomic Analysis Subset* (Subset 3) is designed for use alongside the human genome in experiments focused on genome representative segment selection. It includes the genomes of *Aspergillus nidulans* (Fungi), *Zea mays* (Plantae), and *Dictyostelium discoideum* (Protista). The rationale behind the selection of these species is that, first, the species in this subset are eukaryotes with genome lengths comparable to that of the human genome, enabling similar testing of strategies for representative segment selection. Second, each assembly in this subset is chosen from a different kingdom to evaluate the effectiveness of the analysis and assess the generalizability of findings from the human genome to other eukaryotes. The assemblies in Subset 3 were downloaded from the NCBI database (link in Fig. 1b) on Aug 28, 2024, with additional details and annotations for the maize genome extracted from Hufford et al. [26]. The additional information about the species in this subset is included in Fig. 1b.

### Frequency Chaos Game Representation (FCGR)

Chaos Game Representation (CGR) is a technique to visualize genomic sequences, such as one-dimensional DNA sequences, into two-dimensional visual representations that reflect sequence composition [28]. DNA sequences, composed of four fundamental nucleotide bases—Adenine (A), Cytosine (C), Guanine (G), and Thymine (T)—can be visualized using CGR. To create a CGR image from a sequence, each corner of a square is labeled with one nucleotide. In this analysis, the bottom left corner is labeled A, the top left corner is labeled C, the top right corner is labeled G, and the bottom right corner is labeled T. The center of the square is the starting point, and from there, each nucleotide adds a point to the image, placed halfway between the current point and the corner labeled by that nucleotide [28] (see Fig. 2a). The final CGR image is a two-dimensional plot reflecting the fractal pattern in the DNA sequence composition. Intuitively, for a predetermined *k* value, CGR generates a binary image with size 2^*k*^ × 2^*k*^, where each plotted point corresponds to the presence of a specific *k*-mer in the sequence (see Fig. 2H b). Thus, this technique can be considered both a visualization method and a feature extraction method that encodes the distribution of *k*-mers within a DNA sequence (see Fig. 2H c,d for the examples of CGR images).

**Figure 2:**
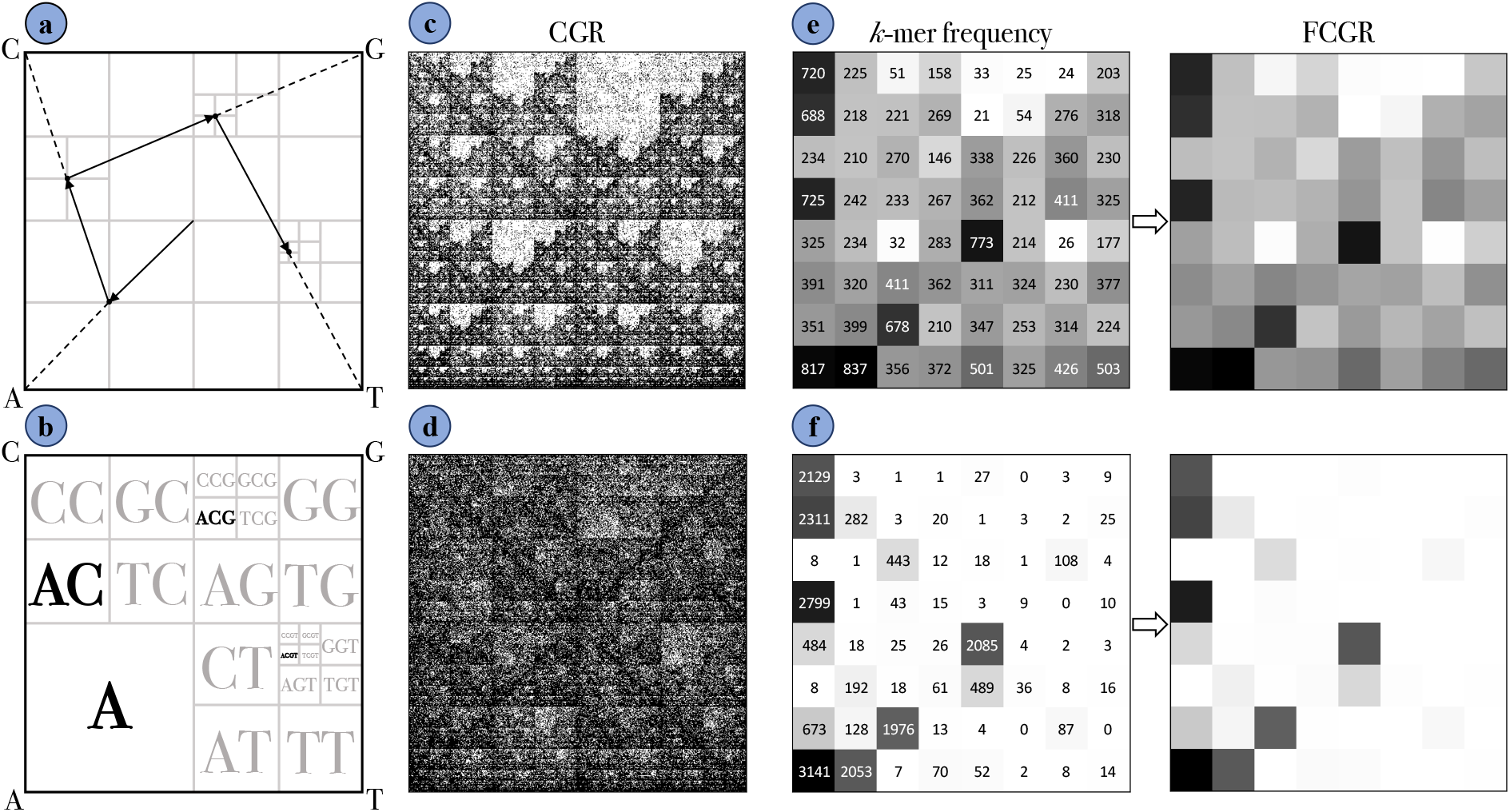
CGR/FCGR image generation. **a**. A schematic of CGR image generation from a DNA sequence. **b**. Mapping of *k*-mers to specific positions in the CGR. **c, d**. Examples of CGR images (512 × 512, *k* = 9) for human (panel c) and maize (panel d). **e, f**. Generating FCGR images by counting *k*-mer frequencies (*k* = 3) for human and maize, respectively. (This figure is adapted from Löchel et al. [46])

The Frequency Chaos Game Representation (FCGR) is an extension of CGR that quantifies the distribution of points generated by CGR [3]. While CGR creates a fractal image of the sequence, FCGR transforms this visual information into a numerical matrix. The unit square is divided into a grid of resolution 2^*k*^ × 2^*k*^, where the intensity of each cell corresponds to the frequency of the *k*-mer within the DNA sequence (see Fig. 2H e,f). The key difference between CGR and FCGR is that CGR displays *k*-mer abundances and biases in *k*-mer composition, while FCGR provides a quantitative representation of the frequency of *k*-mers of a specific value of *k* within the sequence. It is noteworthy to mention that some sequences may contain unknown nucleotides, represented as ‘N.’ During FCGR generation, rather than removing these ‘N’s before extracting the *k*-mers, all *k*-mers are first extracted, and then any containing ‘N’ are discarded. This approach prevents the introduction of unwanted *k*-mers that could result from removing ‘N’ directly from the original sequence.

In generating FCGR images, both the size of *k* and the sequence length influence the visibility of geometric patterns. For a DNA sequence of fixed length, a very small *k* results in low resolution FCGRs, which provide limited detail and make geometric patterns indistinct. Conversely, a very large *k* leads to sparsity in *k*-mer frequencies due to the exponential increase in potential *k*-mers, making the geometric patterns within the FCGR image harder to discern. Similarly, when the image resolution is fixed (i.e., *k* is fixed), shorter sequences contain fewer *k*-mer frequencies, resulting in sparser FCGRs and less discernible geometric patterns. Therefore, an optimal *k* value must be selected for each sequence length to balance high resolution with a sufficient distribution of *k*-mer frequencies, ensuring that the FCGR remains informative and the geometric patterns are clearly visible.

Fig. 2H c depicts the CGR image of the human chromosome 21 and Fig. 2H d shows the CGR of the maize chromosome 8. Their corresponding FCGR representations for *k* = 3 are displayed in Fig. 2H e for human and Fig. 2H f for maize. As seen in the figure, the CGR/FCGR representations differ significantly between the human and maize chromosomes. Generally, FCGRs have been found to be species-specific, such that CGR/FCGR images of DNA sequences from the same genome are quantitatively more similar, while those from different species’ genomes show considerable differences [33]. Therefore, FCGR images have been successfully used in numerous studies for species identification [2, 19, 61, 62], taxonomic classification [1, 43] and phylogenetic clustering [17].

### Distance Measures

Distance measures in this study refer to quantified measures of dissimilarity between FCGR images. Several studies have investigated various methods of comparing two FCGR images [3,31,33,46]. Among the existing methods for measuring the dissimilarity of two FCGR images, eight different distance measures are considered: Normalized Euclidean distance, Cosine distance, Manhattan distance, Structural Dissimilarity Index (DSSIM) [77], Descriptor distance [41], Kolmogorov-Smirnov (K-S) distance [15], Wasserstein distance [50], and Learned Perceptual Image Patch (Dis)Similarity (LPIPS) [83]. Each measure provides a unique way to quantify the dissimilarity between FCGR representations. The Euclidean distance has been widely used in FCGR comparison in different studies [19, 33, 43]. In order to improve the interpretability of the Euclidean distance values, the Normalized Euclidean distance is used, which limits the upper bound of the values. Manhattan, DSSIM, and Descriptor are included as these performed better than the other distances investigated by Karamichalis et al. [33] in a study of intergenomic and intragenomic variation. In addition to well-established distance measures in the literature, four additional distances are incorporated. Cosine distance is included for its interpretability and proven effectiveness in machine learning applications [39, 82]. The K-S statistic and Wasserstein distance are selected as probability-based distance measures, offering different and unique comparison methods with respect to similar studies. Lastly, LPIPS is chosen as a deep learning-based method that combines both structural and perceptual components for measuring dissimilarity. Details on the calculation of each distance measure and the required preprocessing steps are presented in the Appendix A.1.

### Representative Segment Selection Pipeline (RepSeg)

Building on the evidence in this study, demonstrating the pervasive nature of genomic signatures across chromosomes and entire genomes within a species, the Representative Segment Selection Pipeline (RepSeg) is proposed. This pipeline identifies a DNA segment that is significantly shorter than the entire chromosome or genome (e.g., approximately 0.5% of the length of a human chromosome) yet encapsulates the main *k*-mer frequency characteristics of that genome, and closely resembles in this respect most other segments of the same chromosome or genome (hereafter referred to as ‘genome’ for simplicity). The representative segment thus serves as an ideal reference for analyzing intragenomic variations of the genomic signature. Most importantly, the representative segment can act effectively as a proxy of the genome for important applications such as taxonomic classification. This section describes the details of the RepSeg step by step.

The RepSeg begins by dividing a genome into consecutive, non-overlapping segments of equal length. Then, for each segment, it converts the DNA sequence to an FCGR image using a pre-determined *k*-mer value. In the next step, it computes the distance matrix *D* by calculating the distance between all pairs of FCGRs obtained from the segments. The total number of non-overlapping consecutive segments (*S*) determines the dimensions of the resulting *S* × *S* symmetric distance matrix *D*. Finally, it selects the medoid [25] (i.e., the segment that minimizes the average of distances to all other segments in the matrix) and designates it as the representative segment for the genome.

Fig. 3 summarizes all the steps of the RepSeg for a chromosome, including consecutive segment splitting (e.g., size 500 Kbp), FCGR generation (e.g., using *k* = 9), distance matrix calculation, and representative segment selection based on the minimum average distance in the distance matrix. Fig. 3 also includes a Multidimensional Scaling (MDS) representation of the distance matrix *D*, where matrix elements that are close to each other are depicted by closer dots in the MDS plot. The representative segment, highlighted in red, is the center of mass of the points in the MDS representation, as it has the minimum distance to all other points.

**Figure 3:**
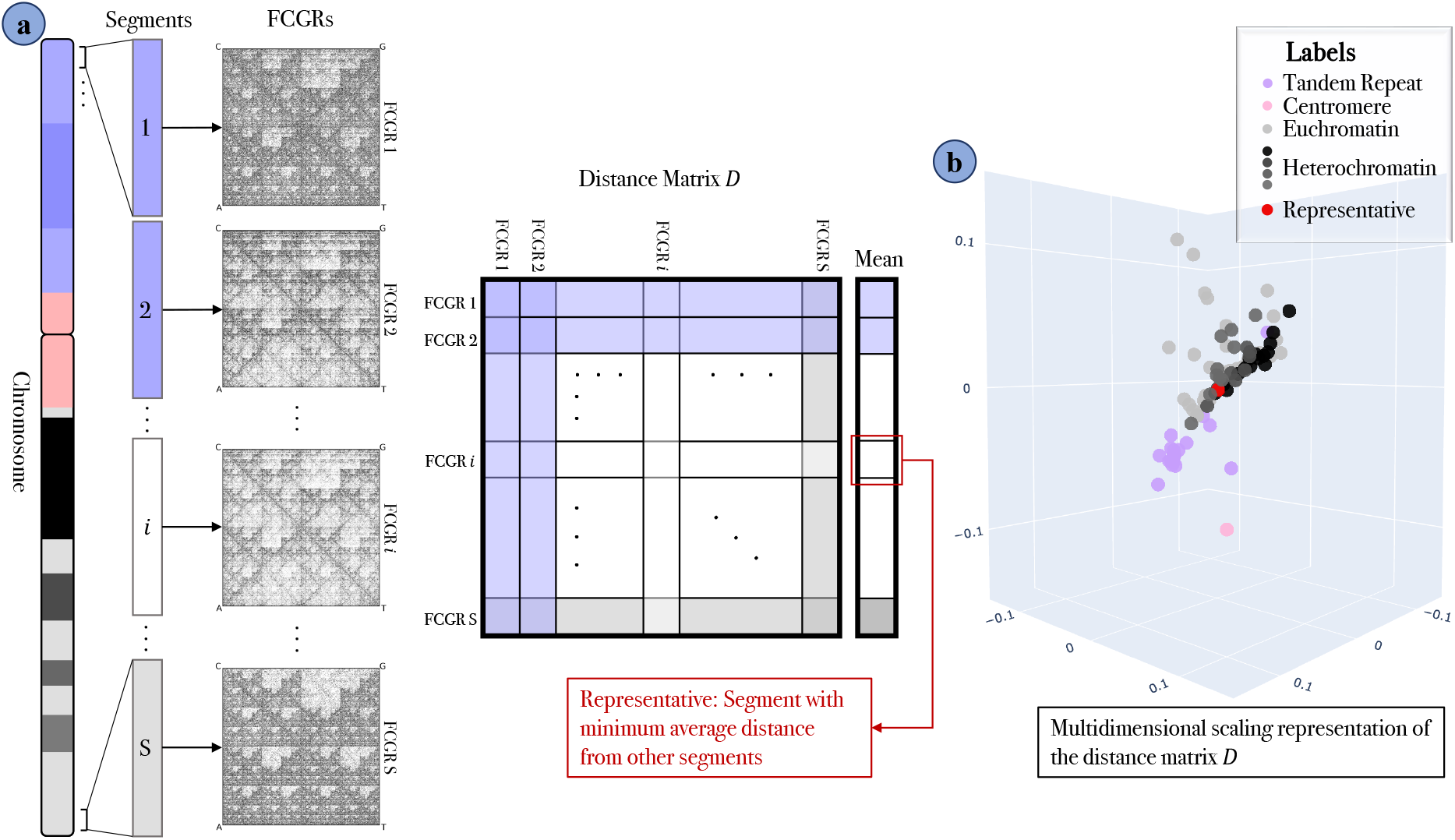
Summary of the Representative Segment Selection Pipeline (RepSeg). **a**. Outlines the steps involved in the pipeline for a chromosome, including chromosome segmentation (e.g., size 500 Kbp), FCGR generation (e.g., using *k* = 9), distance matrix calculation, and representative segment selection. **b**. Multidimensional Scaling (MDS) representation of the distance matrix *D*. The representative segment is highlighted as a red point, representing the center of mass of the MDS plot due to its minimal average distance to other points.

The proposed pipeline identifies the segment with the minimum average distance from all other segments of the genome, making it a strong candidate to serve as the representative segment, as it shares the most similarities with the other segments. However, this pipeline has three intrinsic limitations. First, it is sensitive to both the segment length and the *k* value used in the *k*-mer generation for FCGR (using a different segment length and different *k* can lead to different representative). Second, since the genome is divided into non-overlapping segments, the representative can only be selected from these segments, which means that it is not chosen from among all possible segments of the same length from the genome. Although this issue could be addressed by using overlapping segments, this approach would significantly increase the total number of segments, leading to higher computational and memory costs due to the quadratic time and space complexity of the distance matrix *D*. Finally, computing the distance matrix *D* can be time-consuming for large genomes with a high number of segments, as it requires calculating the distance between all pairs of segments. To address these limitations, the segment length and *k*-value are consistent in our experiments to ensure comparability of results. Subsequently, an approximate pipeline for selecting the representative segment is proposed, which is capable of identifying a representative from any position across the genome, while simultaneously addressing the computational complexity of RepSeg.

### Approximate Representative Segment Selection Pipeline (aRepSeg)

The RepSeg can be computationally expensive due to the quadratic time complexity of distance matrix calculation, which makes its performance highly dependent on the genome length and the chosen segment length. To address this limitation, an alternative pipeline referred to as Approximate Representative Segment Selection Pipeline (aRepSeg) is proposed, which produces a representative segment that approximates the one chosen by RepSeg but with reduced time complexity. For this purpose, instead of dividing the genome into consecutive, non-overlapping segments, aRepSeg randomly selects *n* fixed length segments and computes their pairwise FCGR distances. The number of these random segments affects the time complexity of the algorithm; increasing *n* improves the approximation but also increases the computational cost. The genomic signature, which is pervasive within a species, ensures that the distances between segments of a genome generally exhibit low variability and remain relatively small. However, segments from regions with large tandem repeats can deviate significantly from this pattern, potentially skewing the results. To address this, aRepSeg incorporates a step to identify and exclude these outliers from the randomly selected segments.

More precisely, aRepSeg is an iterative pipeline that starts by initializing the dynamic set *Ŝ* with *n* randomly chosen segments of fixed length. It then creates the distance matrix 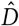 by calculating the pairwise distances between the FCGR images of all segments in the set *Ŝ*. After the calculation of the distance matrix, aRepSeg applies an outlier detection algorithm based on the Interquartile Range (IQR) to identify outlier segments within *Ŝ*. Accordingly, the pipeline removes these outliers from the dynamic set and adds new randomly selected segments, equal in number to the removed ones. The pipeline iteratively recalculates the distance matrix, detects the outliers, and updates the *Ŝ* by replacing the outliers until no further outliers exist in the set. Finally, the segment with the minimum average distance to all others in *Ŝ* is selected as the representative.

To remove outlier segments from *Ŝ*, an algorithm similar to the Interquartile Range (IQR) method is used, which is typically applied to identify and remove outliers. The IQR method works by calculating the interquartile range, which is the difference between the first quartile (Q1) and the third quartile (Q3) of the data [48]. The algorithm then defines a range for typical data by adding 1.5 times the IQR to Q3, known as the upper bound, and subtracting times the IQR from Q1, known as the lower bound. Data points outside this range are considered outliers and are removed, resulting in a cleaner dataset with fewer extreme values. In the pipeline, the IQR method is applied to the average distance of each segment in *Ŝ* from the other segments, which is derived from the distance matrix 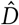. However, aRepSeg modifies this algorithm to only remove values above the upper bound, as small distances are expected, and only high distances pose a problem (they could result from, e.g., one of the segments originating from a tandem repeat region). Any segment detected as an outlier by this modified IQR-based method is removed from the set and replaced with a new random segment chosen from the genome.

To evaluate the effectiveness of aRepSeg and its ability to approximate RepSeg, the representative segments identified by each algorithm are compared. Rather than directly computing the distance between these two representative segments, which would not necessarily be informative, both RepSeg and aRepSeg first identify representative segments for a specific genome. Then, the genome is divided into non-overlapping, continuous segments, each of the same size as the representative segment. Distances are computed between each of these segments and the representative segments selected by RepSeg and aRepSeg, respectively. The Mean Absolute Error (MAE) between the two resulting distance vectors serves as a measure of the approximation error introduced by aRepSeg.

### *CGR-Diff* : A Software for CGR Comparison

To facilitate the experiments on the pervasiveness and variation of genomic signatures, *CGR-Diff*, a novel graphical software tool is developed to extract, visualize, and compare CGRs or FCGRs of different genomic sequences, including chromosomes, genome segments, or any other arbitrary DNA sequence stored in a FASTA file. The tool offers adjustable segment lengths, customizable *k*-mer sizes, and a selection of distance measures for comparison. The software includes a feature allowing users to utilize the assemblies suggested in Fig. 1b or upload their own assemblies for analysis. Additionally, users have the option to select chromosome segments from a list of annotated sequence segments, such as cytobands, if these annotations exist for the selected assembly (cytoband annotations are currently available only for the human genome). The tool also offers the ability to apply the reverse complement to a selected DNA sequence before generating the CGR/FCGR representation. See GitHub for more information regarding how to use the software.

Furthermore, the software includes built-in experiments designed to analyze genomic signature variations both within a species (intragenomic variation) and across different species (intergenomic variation). It also facilitates the replication of the RepSeg and aRepSeg analyses for entire genomes or individual chromosomes from various species. The details of these built-in analyses are as follows:

- *Comparison of consecutive non-overlapping segments for each chromosome or entire genome*: In this experiment, the tool allows users to upload a FASTA file or select the entire genome or specific chromosomes of a species from a predefined list. Users can also specify a segment size and choose a distance measure. The sequence is then divided into consecutive non-overlapping segments of the chosen size. The software then calculates and displays the pairwise distances between neighboring segments, providing a visual representation of these distances on a plot. Additionally, it illustrates the FCGR of each DNA segment, quantifies the distance measures, and visualizes the differences between pairs of FCGRs.
- *Comparison of different segments of a chromosome or genome with a reference sequence*: This experiment is similar to the first, except that the consecutive non-overlapping segments of a chromosome or genome are compared with a chosen, common reference sequence, instead of being compared with their neighboring segments. For the reference sequence, users can choose any segment from the same or different genome with arbitrary start and end points, or select a predefined segment from the list of annotated cytoband sections. The results of these comparisons are visualized in a plot, demonstrating the difference between the chosen reference sequence and each segment. To provide more detailed results, users can select any segment and visually compare its FCGR with that of the reference sequence. The software displays the FCGR of the selected segment, the FCGR of the reference sequence, and the distance between the two, enabling both qualitative and quantitative comparisons.
- *Representative segment selection*: This experiment enables users to run representative selection pipelines on a desired sequence. Users can either upload a FASTA file or select a predefined chromosome or genome from a list. Afterward, they can specify the size of the representative segment, choose the selection pipeline (from RepSeg, aRepSeg, or Random Selection), and select a preferred distance measure. Upon execution, the software identifies and describes the representative segment (including its location in the sequence and its cytoband) and generates a plot showing the distances between different segments along the chromosome or genome and the representative segment. Additionally, it displays the FCGR of the representative segment and each of the segments at each step, along with their differences and distance values.

### Experimental Design

This section provides a detailed account of the experiments conducted in this study, including the hyperparameter values used, such as segment length and the *k* value for FCGR generation. To determine the optimal *k*-mer size, an empirical analysis shows that *k* = 6 offers good visual quality and acceptable quantitative performance for most experiments. However, in Exp 1, where the sequences are larger, a *k* value of 9 is more effective, producing FCGR images with a resolution of 512 × 512 that effectively represents the genomic signatures of full-length chromosomes without causing the FCGRs to fade. Smaller *k* values lacked sufficient resolution, while larger values resulted in faint, less discernible images. For the remaining experiments, where FCGRs are generated from portions of chromosomes rather than entire chromosomes, *k* = 6 is a more appropriate choice, as it was empirically found to deliver better results.

**Exp 1** *Pervasive Nature of Genomic Signatures:* This experiment empirically investigates the pervasive nature of genomic signatures across the genome by visualizing the FCGRs of the full sequences of all 24 human chromosomes and all 10 maize chromosomes using *k*-mers with *k* = 9. This choice of *k* produces highresolution FCGR images, which are empirically verified for representing the genomic signature of the full length of chromosomes. The resolution is particularly critical for detailed visualization, as even the shortest chromosome in these species, human chromosome 21, is approximately 45 Mbp.

**Exp 2** *Distance Selection:* The majority of the experiments are built upon comparing the signature of two DNA sequences using their FCGR images. However, comparing two FCGRs using different distance measures can sometimes lead to non-comparable distance values, as each of them employs a unique method of comparison. Moreover, in FCGR images, it is not only the distribution of *k*-mers that matters; the geometry and visual patterns embedded in the images also contain valuable information that is pervasive and species-specific [53]. Therefore, it is important to analyze the behavior of different distance measures against biological expectations, when comparing FCGR images. For example, if two DNA sequences are phylogenetically similar, the distance value between their FCGRs is expected to be small; otherwise, a larger distance value is expected. To evaluate the effectiveness of the eight distance measures introduced in the section *Methods: Distance Measures* and to determine the optimal one for comparing FCGRs, two experiments are conducted: *Human Intragenomic Distance Analysis*, and *Intergenomic Distance Analysis*.

In the *Human Intragenomic Distance Analysis* experiment, the sequence length varies depending on the region of interest. However, when selecting a random segment from a region, a standard length of 500 Kbp is used. This segment size provides a proper sample size that captures sufficient variation. Since each chromosome spans several Mbp, selecting multiple random 500 Kbp segments ensures diverse sampling and meaningful comparisons. Furthermore, the probability of significant overlap between two randomly selected 500 Kbp segments is very low. For instance, the shortest tandem repeat region within the human chromosome is 10 Mbp, belonging to chromosome 14. Within this region, the chance of observing more than 70% overlap between two randomly selected segments of 500 Kbp is approximately 3%, while the probability of exceeding 50% overlap is only about 5%. These low probabilities ensure that the selected segments are largely independent when the segment size is 500 Kbp, while larger segments would increase the risk of overlap, and smaller segments may fail to capture the genomic signature adequately. Following this rationale, the same 500 Kbp segment length is used in the *Intergenomic Distance Analysis* to maintain consistency.

**Exp 2.1** (*Human Intragenomic Distance Analysis*) focuses on comparing FCGRs within the human genome, leveraging the complexity of this genome and the extensive annotations available for its chromosomes, such as the color-coded regions in the NCBI Data Viewer. In this study, a series of experiments is conducted using various distance measures to compare FCGRs across different human chromosomal regions, including short and long tandem repeats, as well as regions with varying G+C content and CpG composition, such as telomeres, heterochromatin, euchromatin, and the p-arms of acrocentric chromosomes. The effectiveness of these distance measures is evaluated based on their consistency with known differences in sequence composition, such as the occurrence of regional clustering of short and long tandem repeat sequences. This comprehensive intragenomic analysis on the human genome enables the assessment of how well different distance measures align with biological expectations regarding the occurrence and length of tandem repeating sequences. The series of experiments conducted in this study is briefly described below, with detailed explanations provided in the Appendix A.2.

- *Telomere vs. Telomere* aims to determine the distances between the p-arm telomere of chromosome 1 and the p-arm telomeres of other chromosomes. Telomeres, composed of conserved tandem repeats and associated proteins, exhibit similar structure and composition across human chromosomes [44, 67].
- *Heterochromatin vs. Heterochromatin* calculates the average distance between the most condensed heterochromatic region (located distal to the centromere on the p-arm, or, if absent, proximal on the q-arm) and other heterochromatic regions within the same chromosome. The final result is the overall average distance calculated across all chromosomes. Heterochromatin, characterized by high condensation and transcriptional inactivity [60,72], is visually represented by black and three shades of gray in the NCBI Data Viewer, with darker shades indicating higher levels of compaction [22].
- *Heterochromatin vs. Euchromatin* calculates the average distance between the most condensed heterochromatic region (located distal to the centromere on the p-arm or proximal on the q-arm if absent) and four randomly selected euchromatic segments within each chromosome. The final result is the overall average distance computed across all chromosomes. Euchromatin, characterized by less condensed DNA and active gene transcription [8], differs significantly from heterochromatin in structure, function, and genomic composition [8, 52].
- *p-arm vs. q-arm* measures the average distance between 100 randomly selected 500 Kbp segments on the p-arm and q-arm of each acrocentric chromosome (13, 14, 15, 21, 22) as well as the Y chromosome. Acrocentric chromosomes are characterized by a short p-arm, which is enriched with tandem repeat sequences, and a longer q-arm, which contains fewer repeats [49, 69]. These tandem repeat regions consist of long DNA stretches where a sequence is repeated in a head-to-tail manner, varying in repeat unit length, size, and organization [20, 42, 78]. Since the Y chromosome contains repetitive regions in its q-arm [16, 64], it is also included in this experiment.
- *Y q-arm vs. acrocentric chromosome p-arm (Large Tandem Repeat Arrays)* calculates the distance between the tandem repeat arrays on the q-arm of the Y chromosome and those on each acrocentric chromosome, and reports the average. The q-arm of the Y chromosome contains a repetitive region that differs in length and in the nature of its tandem repeat arrays from those on acrocentric chromosomes [64].
- *Arbitrary Sequences* calculates the average intragenomic distance by selecting two random non-overlapping 500 Kbp sequences from a randomly chosen chromosome, computing the distance between their FCGRs, and averaging the results over 100 iterations.

Among the discussed experiments, the *Telomere vs. Telomere* comparison is expected to yield the smallest distance due to the identical repeats in the telomere regions. The *Heterochromatin vs. Heterochromatin* comparison should produce the second smallest distance, and it should be smaller than the *Heterochromatin vs. Euchromatin* comparison. The *p-arm vs. q-arm* experiment, which compares repeat-rich regions to non-repetitive sequences, is expected to result in the largest distance among all tests. The *Large Tandem Repeat Arrays* experiment is expected to show a large distance, but smaller than *p-arm vs. q-arm* since it compares different types of repeats and explores the variation within the repeats, specifically between Y q-arm repeats and acrocentric p-arm repeats. Finally, the *Arbitrary Sequences* test is anticipated to reflect an intermediate intragenomic distance within the human genome, as its large sample size ensures a balanced sampling of diverse combinations of genomic sequences.

**Exp 2.2** (*Intergenomic Distance Analysis*) involves comparing random segments from the human genome with random segments from eight other species listed in Subset 2 of Fig. 1b. Since FCGR images are species-specific, it is hypothesized that phylogenetic distances between the genomes of different species and the human genome will be reflected in their FCGR comparisons, with distances between two random human genome sequences expected to be smaller than those between human and non-human genome sequences. Smaller distances are also anticipated between human and chimpanzee genome sequences compared to those involving human genome and genomes from species in different kingdoms. This evaluation enables the assessment of each distance measure based on how well it aligns with known phylogenetic or evolutionary distances.

To perform this analysis, 100 random segments of length 500 Kbp are selected from the human genome to serve as reference sequences, ensuring that centromeres and large tandem repeat arrays (e.g., the short arms of acrocentric chromosomes and the long arm of the Y chromosome) are excluded. Selections from telomeric regions are not excluded due to their short sequence footprints and minimal impact on the *k*-mer compositions of FCGRs. Then, for each of the nine species (human and eight others), 100 random segments of the same length (500 Kbp) are selected, and pairwise distances between their FCGRs and the reference FCGRs from the human genome are calculated using all eight distance measures. When selecting random segments from the human genome, centromeres and large tandem repeat regions are again avoided. Finally, these distances are compared across all nine species using boxplots and the Wilcoxon signed-rank test [63, 80], with expectations based on known phylogenetic relationships. The Wilcoxon signed-rank test is applied because the same reference human FCGRs are used to compute both sets of distances, leading to dependent matched samples.

**Exp 3** *Intragenomic Variation*: A series of experiments is conducted to evaluate the RepSeg and aRepSeg methods. The methods that are designed to select representative genomic segments for various species. These representatives serve multiple purposes, including exploring variation in genomic signatures along chromosomes or entire genomes and supporting downstream tasks, such as the species classification in Exp 4. Experiments in this section, focus on the human and three species listed in Subset 3 of Fig. 1b. For human and maize, which have long genomes (approximately 3 billion bp and 2 billion bp, respectively) and chromosomes ranging from 45 Mbp (e.g., human chromosome 21) to 300 Mbp (e.g., maize chromosome 1), a representative segment is selected for individual chromosomes to maintain their distinct genomic information. For species with smaller genomes, such as *Aspergillus nidulans* and *Dictyostelium discoideum*, all chromosomes are concatenated with ‘N’ to prevent unwanted *k*-mers, and a representative segment is selected for the entire genome. In all cases, a segment length of 500 Kbp is used, as it is sufficiently long to preserve genomic signatures while remaining a small fraction of the total genome length (e.g., less than 1% of the shortest human chromosome). Furthermore, for comparing FCGRs in these experiments, the DSSIM distance measure is utilized, as it demonstrates the best compatibility with the biological expectations discussed in the Distance Analysis experiment (see *Results* section).

**Exp 3.1** (*Representative Segment Selection*) applies the RepSeg to the human genome and the genome of each species listed in Subset 3 of Fig. 1b. This pipeline finds a representative segment for each individual chromosome in human and maize, and for the entire genome in *Aspergillus nidulans* and *Dictyostelium discoideum*. Using the representative segment as a reference, the experiment calculates and plots the distances between each consecutive segment (of the same length as the representative segment, i.e., 500 Kbp) and the representative segment. This analysis highlights the prevalence or variation of the genomic signature within a single chromosome or the entire genome.

**Exp 3.2** (*Approximate Representative Segment Selection*) repeats the previous experiment on the human genome and the genomes of all the species of Subset 3 in Fig. 1b, but uses the aRepSeg to determine the representative segment. The aRepSeg utilizes a set of random segments *Ŝ*, and in this experiment, the size of *Ŝ* is set to 30. This size is determined through an empirical experiment (see Supplementary Table S4).

**Exp 3.3** (*Unlikely Representative Segment from Tandem Repeat Regions*) is an experiment similar to the previous ones but designed to demonstrate the impact of selecting a random segment from regions with large and small tandem repeats, such as the centromere, as the representative segment. This experiment is applied only to human chromosomes, as it is the only genome from Fig. 1b that contains annotations for cytobands and repeat regions.

**Exp 4** *Taxonomic Classification*: To evaluate the utility of the representative segment chosen by RepSeg or aRepSeg in downstream tasks, a simple taxonomic classification experiment is conducted using a one-nearest-neighbor (1-NN) classifier. This experiment is applied to all the species listed in Fig. 1b, except for *Paramecium caudatum*, whose genome is available only at the scaffold level, making it unsuitable for consistent segmentation or classification. For this experiment, the segment size is reduced from 500 Kbp, as used in previous experiments, to 200 Kbp. This adjustment accommodates the relatively short genomes of some species in the dataset, where extracting 100 random 500 Kbp segments per species would be impractical.

**Train Data**: The training dataset consists of one representative for each species, which is selected under two scenarios: (1) using the pipelines (RepSeg or aRepSeg) or (2) choosing a random segment from a chromosome or genome as the representative. For genomes shorter than 100 Mbp, in the first scenario, the representative is selected by applying RepSeg or aRepSeg to the entire genome, whereas in the second scenario, the genome is divided into consecutive segments of 200 Kbp, and one segment is randomly chosen as the representative. For genomes longer than 100 Mbp, the representative in the first scenario is determined as the final representative of the representatives of individual chromosomes, where the final representative is the segment with the minimum average distance to all other chromosome-level representatives. In the second scenario, a random chromosome is first selected, divided into consecutive segments of 200 Kbp, and a random segment from these is chosen as the representative.

**Test Data**: The test dataset consists of 100 random segments of 200 Kbp each, extracted from the entire genome for species with genomes under 100 Mbp and from random chromosomes for species with genomes exceeding 100 Mbp. This results in a total of 1,100 test samples, representing 11 species.

**Classification Approach**: The classifier predicts the label for each test sample by comparing the DSSIM distance between the FCGR of the test sample and the FCGRs of the 11 representative segments in the training set. The DSSIM distance measure is used for comparing FCGRs, as described in Exp 3.

**Evaluation**: The two scenarios are compared by computing their classification accuracy. To avoid bias towards tandem repeat regions, for the random selection scenario, the average accuracy over 50 repetitions is reported. This comparison assesses the effectiveness of the pipelines (RepSeg or aRepSeg) in generating representative segments for downstream tasks.

## Results

### *CGR-Diff* Software Tool for Assessing Intragenomic Variations in the Genomic Signature

Fig. 4 demonstrates the capabilities of the software tool by presenting an example from its first built-in experiment, *Comparison of consecutive non-overlapping segments for each chromosome or entire genome*. The primary objective of this experiment is to evaluate intragenomic variation in the genomic signature and to quantitatively compare genomic signatures along the chromosome. In this example, human chromosome 1, the longest chromosome in the genome, is analyzed using a segment size of 500 Kbp, *k* = 6 for FCGR generation, and DSSIM (as defined in *Methods: Distance Measures*) as the distance measure to capture variation along the chromosome.

**Figure 4:**
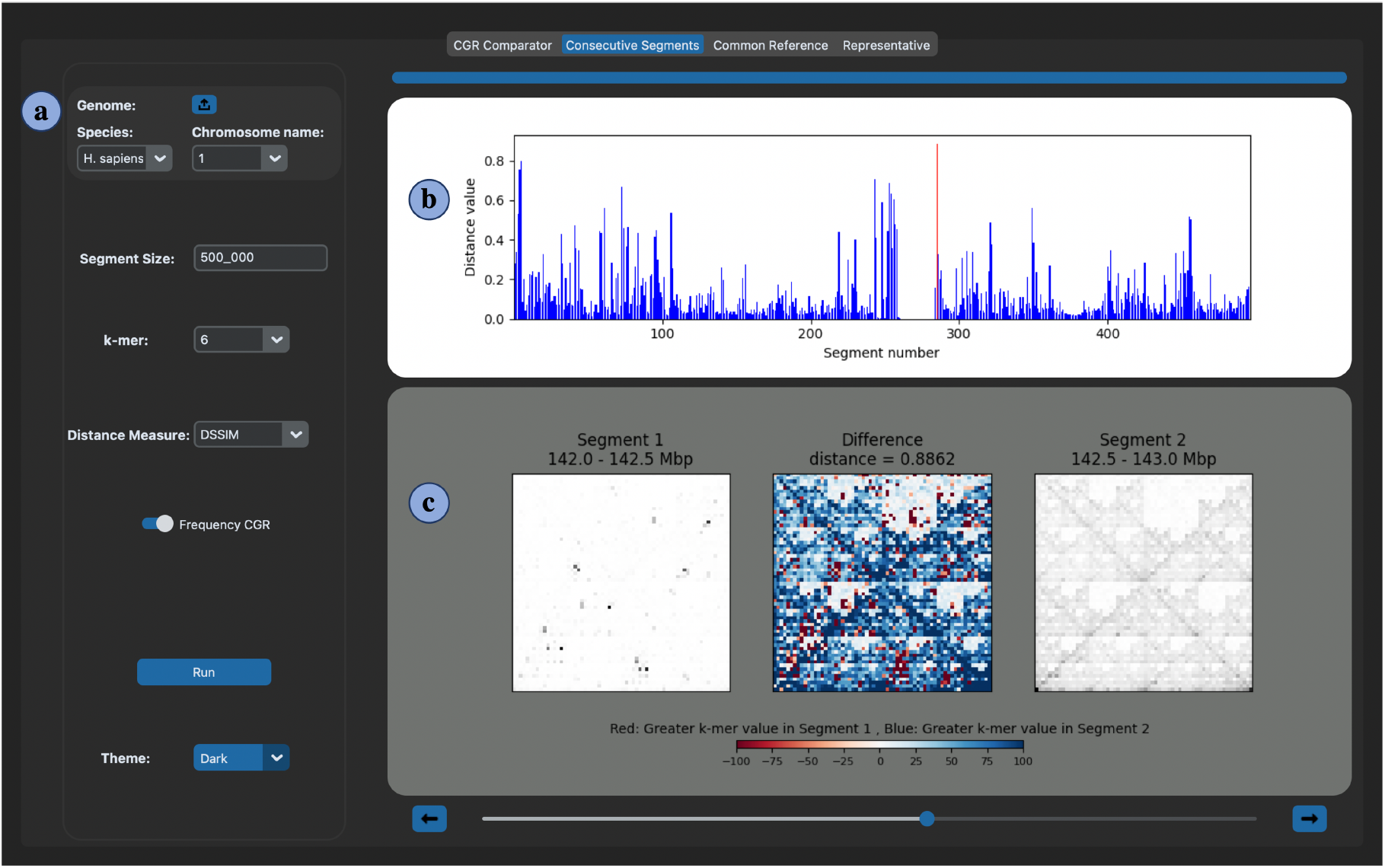
A screenshot of the *consecutive non-overlapping segments* experiment of the *CGR-Diff* software. **a**. Control panel showing the parameters of the experiment, including *k*, segment size, and distance measure. **b**. Plot displaying the distances between FCGRs of consecutive segments across the first human chromosome (using *k* = 6, segment size = 500 Kbp, and distance measure = DSSIM). The red bar indicates the maximum distance (0.88) at the boundary between a tandem repeat region (q12) and a euchromatic region (q21.1). **c**. FCGRs correspond to two consecutive segments associated with the maximum distance, with their positions on the chromosome mentioned at the top of the images. The left and right images show the individual FCGRs, and the center shows their pixel-wise difference, highlighting shifts in *k*-mer composition.

Fig. 4H b displays a feature of the software that visualizes the DSSIM distance between each pair of consecutive genomic segments. As seen in this figure, in the case of human chromosome 1, most of these distance values are low (with an average of approximately 0.11), indicating a consistent genomic signature between neighboring segments. However, several high distance values are observed to occur at specific locations, particularly at cytoband boundaries, reflecting significant changes in *k*-mer composition. The highest observed distance in this experiment is 0.88 (highlighted in red), which is a significant value given that DSSIM ranges from 0 to 1 when comparing FCGR images. This peak occurs at the boundary between the q12 and q21.1 cytobands of chromosome 1, where q12 is associated with tandem repeats, while q21.1 corresponds to a euchromatin region. Before this peak, there is a portion of the plot where the DSSIM distances are close to zero. This low variation occurs within the tandem repeat region (q12), where the sequences of consecutive segments are highly similar, resulting in similar FCGR patterns and low DSSIM distance values between them.

Fig. 4H c displays a feature of the software that provides a detailed visualization of two selected consecutive segments and their comparison. In this example, the two selected consecutive segments are those whose distance is the peak distance value in Fig. 4H b (segments 285 and 286, together spanning 142.0–143.0 Mbp). The left and right images display the FCGRs of the individual segments, while the center matrix shows the pixel-wise difference between them, highlighting which *k*-mers are more abundant in each segment. The observed difference in FCGR patterns reveals a shift in sequence composition that aligns with the transition from a tandem repeat-rich region (Segment 1) to a euchromatic region (Segment 2).

### Demonstrating the Pervasive Nature of Genomic Signatures (Exp 1)

The FCGRs for the complete sequences of all 24 human chromosomes (see Fig. 5) and all 10 maize chromosomes (see Supplementary Fig. S1) are generated using *k* = 9. The FCGRs of human chromosomes exhibit similar geometric patterns across all chromosomes, which are distinctly different from those observed in maize. Similarly, the FCGRs of maize chromosomes are consistent among themselves. These observations support the assumption that genomic signatures are both species-specific and pervasive across the entire genome of a species.

**Figure 5:**
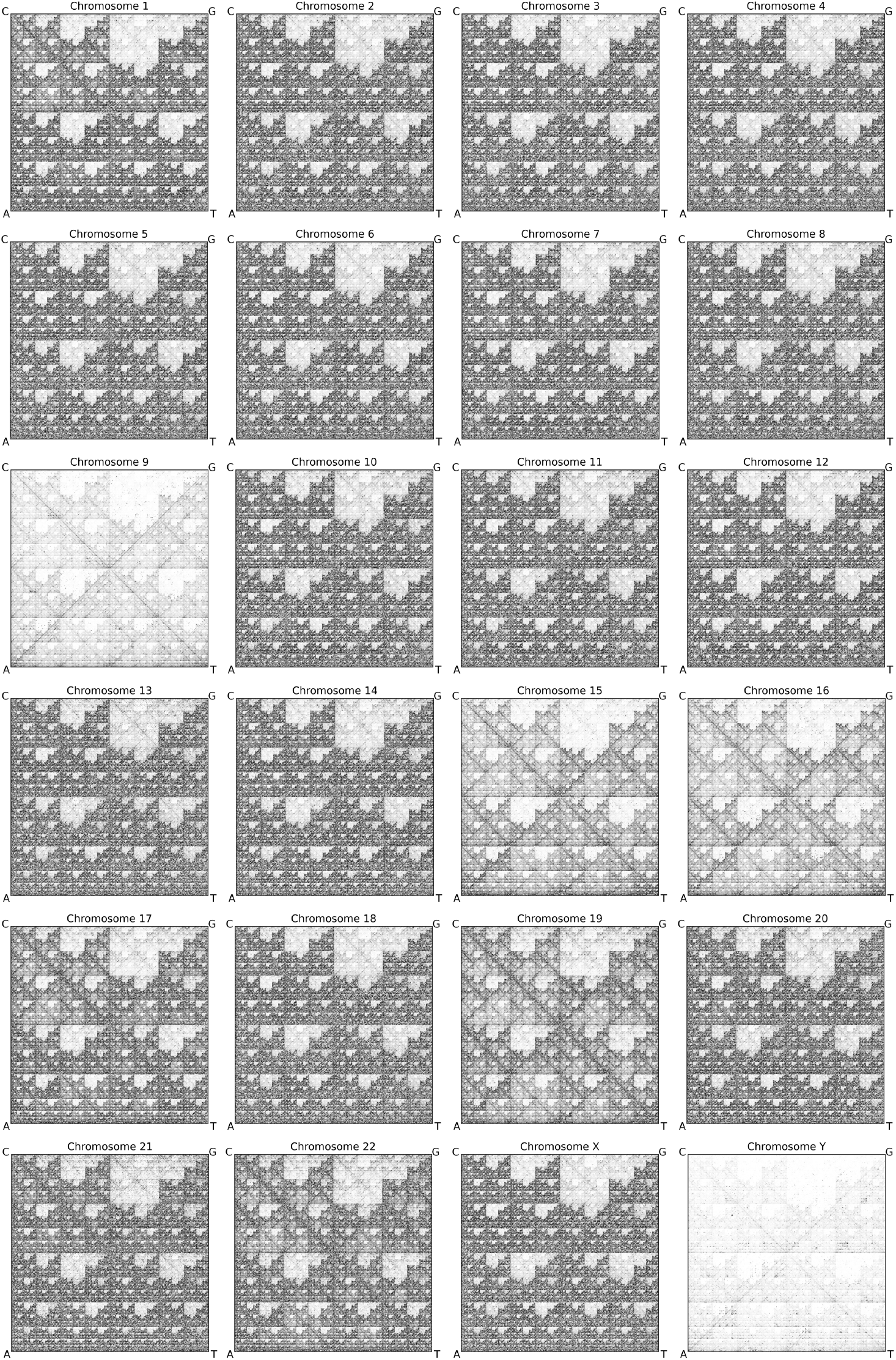
Analyses of the Human Genome Genetic Signature. Each image represents the FCGR of a complete human chromosome, constructed using *k* = 9. The overall structure reveals a preserved genomic signature across chromosomes, while variations in intensity indicate differences in *k*-mer distribution. Specifically, certain chromosomes, such as chromosome 9, 15, 16, and Y, appear lighter, consistent with the presence of regions with known high *k*-mer repetition.

Despite the overall consistency of FCGR patterns across all human chromosomes, chromosomes 9, 15, 16, and Y exhibit notable deviations in FCGR intensity, as shown in Fig. 5. These variations are not due to chromosome length, but rather to the high relative frequencies of some specific *k*-mers (9-mers in Fig. 5), which result in lower counts of the other *k*-mers (lighter other regions in the FCGR image). Specifically, the relative frequencies of AAGGTAAGG (chromosome 9, 0.75%), CTTACCTTA (chromosome 15, 0.34%), TTACCTTAG (chromosome 16, 0.36%), and TAAGGTAAG (chromosome Y, 1.00%) are significantly higher than the relative frequency of 9-mers in the other chromosomes (average 0.13%). These observed high frequencies of specific 9-mers are likely the result of extensive repetitive regions within chromosomes 9, 15, 16 and Y, and are absent in the other chromosomes (see Supplementary Table S1). For example, chromosome 9 contains the largest block of heterochromatin among human chromosomes and exhibits numerous repetitive regions within the centromere and large heterochromatic regions [27]. Similarly, chromosomes 15 and 16 have some of the highest levels of segmental duplications in the human genome [68, 85]. Finally, the human Y chromosome is substantially different from all other chromosomes, as it is densely packed with repeats, such that almost any sequence from the Y chromosome either repeats internally or has a near-identical copy elsewhere on this chromosome [64, 66].

A similar phenomenon is observed in the maize chromosomes, with the FCGR images of chromosomes 2 and 4 displaying reduced overall pixel intensities. These lighter FCGRs are due to the high relative frequencies of specific *k*-mers (see Supplementary Table S2). Specifically, the relative frequencies of TCATCATCA (chromosome 2, 0.10%) and TGATGATGA (chromosome 4, 0.05%), are higher than the relative frequency of 9-mers in the other chromosomes (average 0.02%).

### Determining the Optimal Distance Measure (Exp 2)

#### Human Intragenomic Comparison (Exp 2.1)

The performance of the eight distance measures in the human intragenomic distance analysis experiment is shown in Fig. 6. Considering the identical sequence repeats in telomeres, the *Telomere vs. Telomere* comparison is expected to yield the smallest distance. As seen in the figure, this expectation is met for all distance measures except Normalized Euclidean, Cosine, and Wasserstein distances. Additionally, the expectation that the *Heterochromatin vs. Euchromatin* comparison would result in a larger distance than the *Heterochromatin vs. Heterochromatin* comparison is confirmed by all distance measures. The *p-arm vs. q-arm* experiment in different chromosomes is expected to produce the largest distance among all tests, as it compares repeat-rich regions to non-repetitive sequences. However, as shown in Fig. 6, the Normalized Euclidean, Cosine, K-S, and Wasserstein distance measures fail to reflect this expected result. Finally, in the *Large Tandem Repeat Arrays* experiment, a large distance is expected, greater than in the *Arbitrary Sequences* comparison but smaller than in the *p-arm vs. q-arm* comparison, since it compares different types of large tandem repeat arrays. However, the Normalized Euclidean, Cosine, Manhattan, K-S, and Wasserstein distance measures fail to meet this expectation. Overall, among all the distance measures, Descriptor, DSSIM, and LPIPS performed as expected in Exp 2.1.

**Figure 6:**
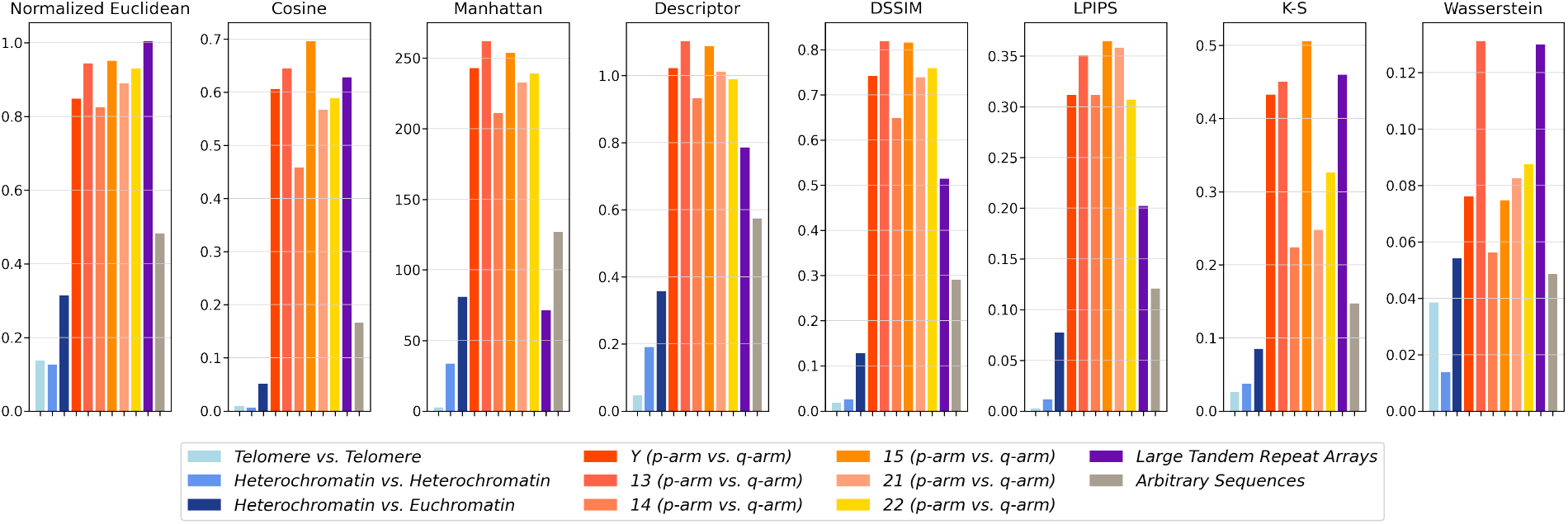
Intragenomic Distance Analysis (Exp 2.1). Each bar plot shows the performance of a distance measure across the different experiments in Exp 2.1. Three shades of blue represent the *Telomere vs. Telomere, Heterochromatin vs. Heterochromatin*, and *Heterochromatin vs. Euchromatin* experiments, which are expected to yield relatively small distance values compared to the other experiments (the lighter the shade, the smaller the expected distance value). Six warm colors correspond to the *p-arm vs. q-arm* experiment in different chromosomes (Y, 13, 14, 15, 21, and 22), where distance values are expected to be the largest among all experiments. Purple bars represent the *Large Tandem Repeat Arrays* experiment, which is expected to have large distance values, though smaller than those in the *p-arm vs. q-arm* experiment. Finally, gray bars indicate the *Arbitrary Sequences* experiment, which is expected to produce intermediate distance values. Among all distance measures, Descriptor, DSSIM, and LPIPS align best with biological expectations.

#### Intergenomic Comparisons (Exp 2.2)

Fig. 7 presents boxplots illustrating the distribution of intergenomic distance values across species with known phylogenetic distances from the human genome, using the eight proposed distance measures. Each boxplot represents the distribution of pairwise distances between 100 randomly selected human segments (referred to as human reference segments) and 100 randomly selected segments from the corresponding species. In the arrangement of the boxplots, the first plot, positioned on the far left, corresponds to comparisons between human reference segments and randomly selected human genome segments (human-human). Progressing to the right, the subsequent boxplots represent increasing phylogenetic distances, comparing human reference segments to segments from increasingly distantly related species. This arrangement illustrates the expected trend of increasing intergenomic distances as we move from closely related species (e.g., *Pan troglodytes* (chimpanzee) and *Mus musculus* (mouse)) to those less related, and further to species from different kingdoms.

**Figure 7:**
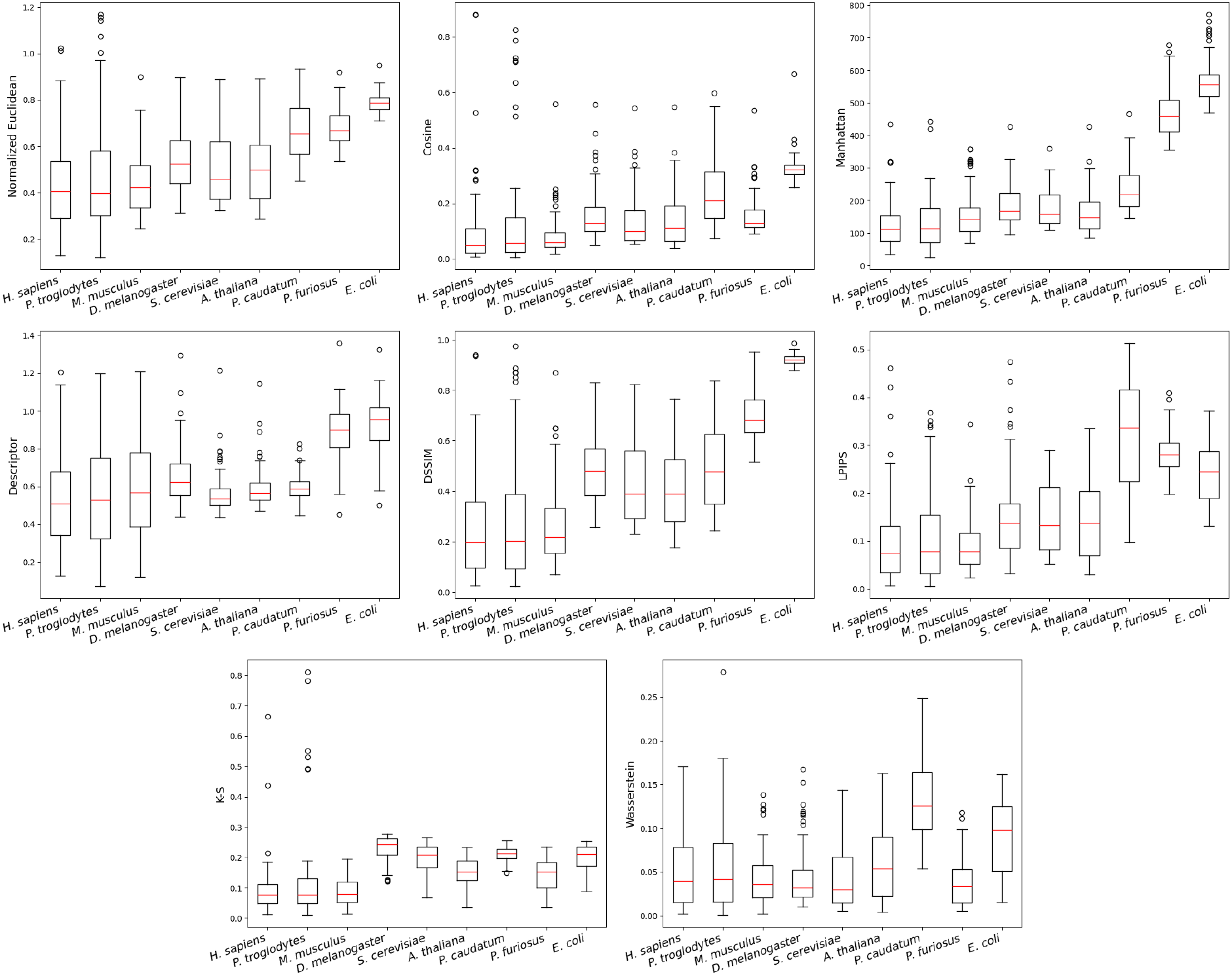
Intergenomic Distance Analysis (Exp 2.2). Boxplots of eight distance measures showing the distance distribution between FCGR segments from 100 randomly selected human reference segments and 100 randomly selected segments from corresponding species. The first boxplot in each plot represents comparisons within the human genome, while the rest show comparisons between human reference segments and randomly selected segments from other species.

Based on biological hypotheses, an increase in distance values is expected from left to right along the horizontal axis in each plot, reflecting greater divergence from the human genome. Smaller distances are anticipated for comparisons between human reference segments and randomly selected human genome segments (human-human) compared to distances between human reference segments and segments from other species. Furthermore, no significant difference (i.e., *p*-value greater than 0.05) is expected between the human-human distances and the distances between human reference segments and segments from closely related mammals, such as *Pan troglodytes* (chimpanzee) and *Mus musculus* (mouse). In contrast, significant differences are expected when comparing the human-human distances to the distances between human reference segments and segments from species in different kingdoms.

Moreover, the variability in distance distributions when comparing each species to human reference segments reflects the intragenomic variation within that species. This variation is due to known intragenomic sequence composition, particularly in relation to large tandem repeat regions. Therefore, the variability of each boxplot can also be assessed based on biological predictions. For instance, the genome composition of vertebrates leads to an expectation of greater variability compared to bacterial genomes. Notably, the chimpanzee genome is predicted to exhibit a high variation, highlighting its abundant intragenomic differences.

According to Fig. 7 and the *p*-values from the Wilcoxon signed-rank test in Supplementary Table S3, no statistically significant differences are observed between the human-human distances and the human-*P. troglodytes* (chimpanzee) distances across the eight distance measures. However, when comparing the human-human distance values with those of human-*M. musculus*, the Manhattan, Descriptor, and Wasserstein distance measures yield statistically significant differences, with *p*-values of 0.0009, 0.0348, and 0.0415, respectively. Therefore, these three distance measures, do not support the expectation of no significant difference among mammals. Additionally, when comparing the human-human distances with the distances between human and more distantly related species, all distance measures show statistically significant differences, except for Wasserstein distance in the comparison between human-human and human-*A. thaliana* (*p* = 0.0684). Among the remaining distance measures, Normalized Euclidean, Cosine, and DSSIM most accurately reflect phylogenetic distances and align with the expected variation in vertebrates.

#### Conclusion of Exp 2: DSSIM Confirmed as Preferred Distance Measure

Considering the results of Exp 2.1 and Exp 2.2, DSSIM is the preferred distance measure for comparing FCGRs and is used in subsequent experiments. The consistent performance of DSSIM is further supported by the Wilcoxon signed-rank test, where the *p*-value for the comparison between the human-human distances and those of human-*P. troglodytes* is 0.178, between human-human and human-*M. musculus* is 0.096, and for all other species, the *p*-value is less than 10^−11^. The DSSIM distance measure not only meets all biological expectations but also, compared to distance measures like LPIPS (as defined in *Methods: Distance Measures*), has lower computational complexity. Additionally, it has the advantage of boundedness, with a practical range of [0, 1], which makes the interpretation of numerical results easier.

### Evidence of Low Intragenomic Variability of the Genomic Signature (Exp 3)

To quantitatively measure the intragenomic variation of genomic signatures within each chromosome of humans and maize, as well as across the entire genomes of *Aspergillus nidulans* and *Dictyostelium discoideum*, the RepSeg and aRepSeg are applied as described in Experiments 3.1 and 3.2. These pipelines extract representative segments, and the distances between consecutive segments (each 500 Kbp in length) and the representative segments are calculated.

Fig. 8 presents the results of these experiments for human chromosomes. The line plots show the DSSIM distances of segments relative to their representative segments. In the red lines, the representative segment is selected using the RepSeg method, which follows a deterministic pipeline using all non-overlapping consecutive segments. The blue lines show the distances of the segments from the representative segment selected using the aRepSeg method, which employs an approximation pipeline based on a random sample of segments. Finally, the black lines illustrate the distances of consecutive segments from a randomly selected segment within regions of high repetitive sequences, as described in Exp 3.3. The Fig. 8 also includes ideograms of human chromosomes (adapted from the NCBI Genome Data Viewer [54]).

**Figure 8:**
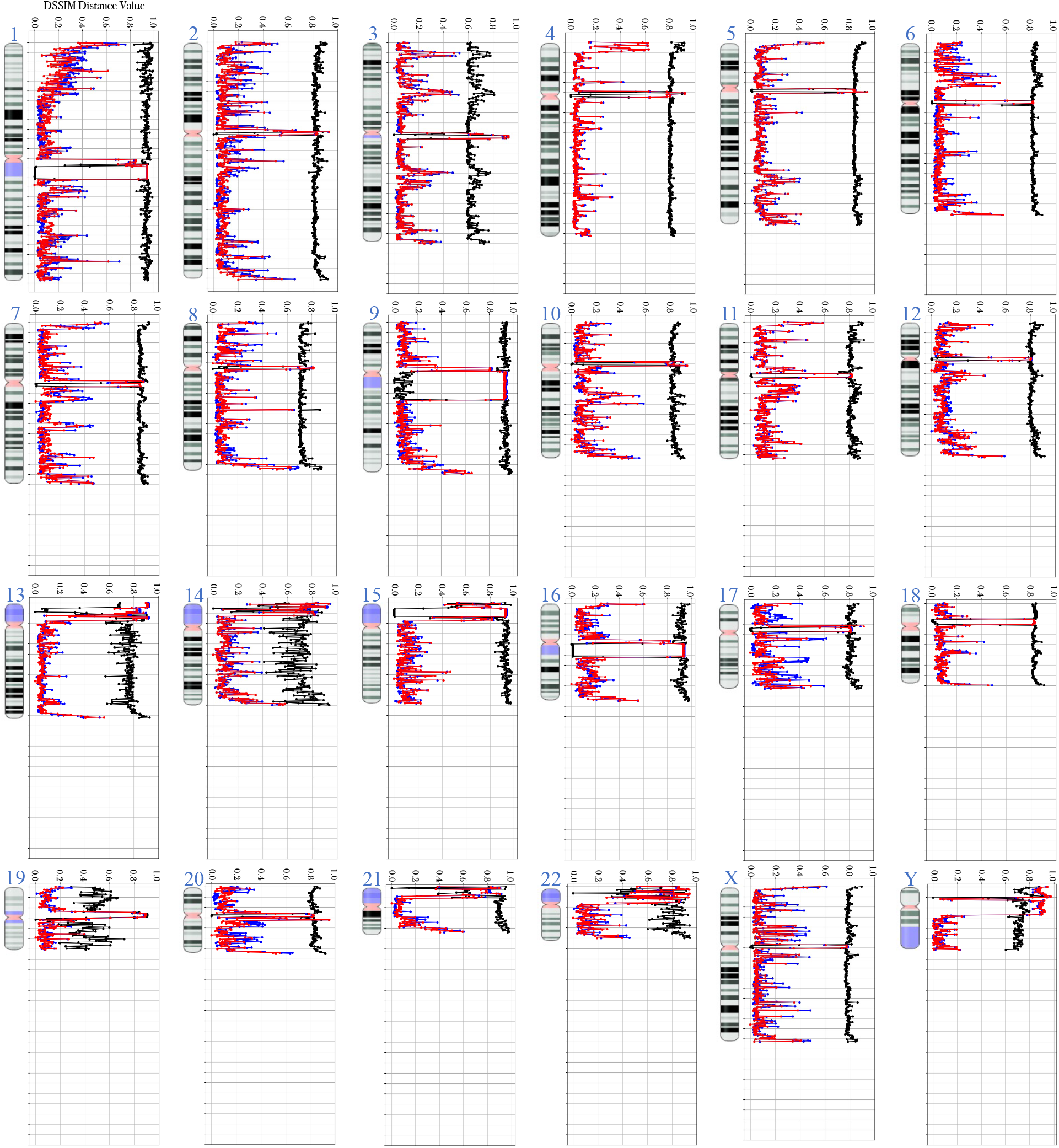
DSSIM distances of consecutive segments from the representative segment across all human chromosomes. The red, blue, and black lines correspond to three experiments (Exp 3.1, Exp 3.2, and Exp 3.3), with each point along the line representing the calculated distance between a 500 Kbp genomic segment and a representative segment selected by the proposed pipeline. The horizontal axis shows the DSSIM distance values ranging from 0 to 1 (with increments of 0.2), where higher values indicate greater dissimilarity. The vertical axis is divided into intervals of 20, corresponding to sequential 500 Kbp segments along each chromosome. Chromosome ideograms to the left of each plot are adapted from the NCBI Genome Data Viewer [54] and display key structural regions such as heterochromatin (regions colored in black and three shades of gray), euchromatin (white regions), centromeres (pink regions), and large tandem repeat arrays (purple regions).

According to the red lines in Fig. 8, among all human chromosomes, the majority of segments (83.32%) exhibit a DSSIM distance of less than 0.24 from the representative segment suggested by RepSeg. Overall, only a small fraction of segments (16.67%) display outlier behavior, with FCGR patterns that differ from the majority and are associated with highly repetitive sequences in the chromosome. The similarity of FCGR patterns across most segments confirms the pervasiveness of genomic signatures within the chromosomes. This finding aligns with the results of Experiment 1, which provided evidence for the pervasiveness of the genomic signature across different chromosomes, further supporting their pervasiveness within individual chromosomes as well. Consequently, it is reasonable to propose a shorter representative segment for a chromosome with millions of base pairs that effectively captures this signature.

As the genomic signature is consistently repeated along the chromosome, one may assume that a randomly selected segment from the chromosome can also serve as the representative. However, selecting a purely random segment as the representative is not advisable, as a randomly chosen segment from the outlier segments is an unlikely representative and would fail to accurately encapsulate the genomic signature (see the substantial difference between the black and red lines in Fig. 8 with an average MAE of 0.61 across all chromosomes). On the other hand, the close resemblance (average MAE of 0.02 across all chromosomes) between the blue line (aRepSeg) and red line (RepSeg) further validates the effectiveness of aRepSeg in accurately capturing the genomic signature while reducing the computational cost of RepSeg.

Supplementary Table S4 shows the effect of the hyperparameter *n* (the size of the set *Ŝ* in aRepSeg) on runtime improvement and MAE between aRepSeg and RepSeg. In this table, an *n* value of 1 corresponds to random representative selection. Intuitively, *n* determines the size of a dynamic set from which the representative segment is selected—only if the set contains no outliers. A larger *n* increases the chances of including a high-quality representative segment; however, larger values of *n* also increase time complexity, thereby reducing the time-saving advantage of aRepSeg. Therefore, identifying an optimal *n* is essential to balance both time efficiency and effective representative selection. Computational analysis, which explored values of *n* between 1 and 50, empirically determined that *n* = 30 achieves an MAE of 0.027 on human chromosome 1, which is a negligible error within the range of DSSIM distances and corresponds to approximately a 20x improvement in computational time for representative selection in this chromosome.

In summary, the study confirms that a representative segment of 500 Kbp, as suggested by both RepSeg and aRepSeg, effectively captures the genomic signature within chromosomes. This representative facilitates intragenomic variation analysis, and quantitative exploration of the pervasiveness of genomic signatures along individual chromosomes of the human genome, while highlighting outlier regions that show abnormal genomic signatures. Moreover, setting the hyperparameter *n* to an optimal value of 30 for aRepSeg keeps a balance between precision and computational efficiency.

To demonstrate the generalizability of RepSeg and aRepSeg beyond the human genome, these pipelines are applied to ten maize chromosomes (see Supplementary Fig. S2) as well as the entire genomes of *Aspergillus nidulans* and *Dictyostelium discoideum* (see Supplementary Fig. S3). The results from these species are consistent with those observed in Fig. 8, further supporting the pervasive nature of genomic signatures across both chromosomes and entire genomes. Some variations are observed, primarily due to the presence of tandem repeat regions, which is a pattern also seen in the human genome.

Across all maize chromosomes, the majority of segments (93.23%) exhibit a DSSIM distance of less than 0.24 from the representative segment selected by RepSeg. Furthermore, aRepSeg closely replicates RepSeg across all chromosomes, with an average MAE of 0.02. Also, similar to the human genome, DSSIM distances in the maize genome tend to increase in segments associated with centromeres and large tandem repeat regions.

The assembly of *Aspergillus nidulans* consists of eight chromosomes, which are concatenated to form the full genome sequence. Since the individual chromosomes have an average length of 3.7 Mbp, the 500 Kbp representative selection pipeline is applied to the concatenated chromosomes rather than to individual ones. Similarly, *Dictyostelium discoideum* has six chromosomes with an average length of 5.6 Mbp, and the same 500 Kbp representative segment selection pipeline is applied to the whole genome obtained by concatenation of all chromosomes. Supplementary Fig. S3 illustrates the DSSIM distance of consecutive segments from the representative segment selected by RepSeg (red line) and aRepSeg (blue line) for each species. Compared to the human and maize genomes, these two eukaryotes exhibit a more uniform distance from the representative segment, further supporting the pervasiveness of genomic signatures within the genome of a species. In *Aspergillus nidulans*, the average DSSIM distance from the representative segment selected by RepSeg is 0.13, with a standard deviation of 0.03, indicating low intragenomic variation. Also, aRepSeg closely aligns with RepSeg, with an MAE of 0.01. A similar trend is observed in *Dictyostelium discoideum*, where the average DSSIM distance is 0.004 (standard deviation: 0.006), and aRepSeg deviates from RepSeg with an MAE of 0.0003.

### Effectiveness of Genomic Signatures for Alignment-Free Taxonomic Classification (Exp 4)

To demonstrate the effectiveness of the representative segment selection pipelines, a simple taxonomic classification task is performed using 1,100 randomly selected segments from the species listed in Fig. 1b. First, a training set is constructed by selecting the representative segment identified by either RepSeg or aRepSeg for each species. Then, test samples are classified based on their DSSIM distance to the corresponding representative segment, achieving an accuracy of 84.91% when using RepSeg and 84.45% when using aRepSeg to select the representative segments.

Furthermore, to assess the significance of the representative segments selected by the pipelines, the classification task is repeated using a randomly chosen segment from each species as the reference instead of the representative segment suggested by the pipelines. This substitution results in a drop in average classification accuracy to 77.63% over 50 runs. This reduction in accuracy highlights the effectiveness of representative segment selection for downstream applications such as taxonomic classification. Additionally, it suggests that while genomic signatures are pervasive across the genome of a species, they are not entirely uniform, likely due to the presence of repetitive regions.

Due to the high similarity in the genomic signatures of humans and chimpanzees, most misclassifications occur between these two species (see confusion matrices in Supplementary Fig. S4). To address this, we repeat the experiment excluding chimpanzees from the dataset and achieve an accuracy of 91.7% using RepSeg and 91.4% using aRepSeg. In comparison, the average accuracy using random segments as representative segments across 50 runs is 84.36%. While the accuracy improves in all scenarios, the difference between the accuracy using the pipeline-selected representative and the accuracy using a random representative remains the same.

## Discussion and Conclusions

This study investigates the intragenomic variation of genomic signatures through the analysis of *k*-mer distributions in FCGR images and proposes effective methods to select a representative segment to serve as a proxy for the whole genome for taxonomic classification or other applications.

Overall, our findings indicate that the *k*-mer distribution reflected in FCGR images is preserved throughout the genome of the studied species, with some exceptions in repetitive regions. This counterintuitive pervasiveness of the genomic signature suggests that *k*-mer compositions within DNA sequences are shaped by fundamental patterns that tend to persist over long periods of time [17]. This being said, our analysis revealed some notable instances of FCGR patterns that deviate from the dominant genomic signature, suggesting that the pervasiveness of the genomic signature has exceptions. These exceptions, though limited in scope, may signal important biological events, and examining how and why genomic signature patterns vary in certain regions could provide valuable insights into evolutionary processes, adaptations, the detection of genomic islands, horizontal gene transfer events, and structural annotation. For instance, we hypothesize that genomic islands, which have acquired genes from other organisms, could exhibit abnormal genomic signatures compared to the host genome that might be identified through the intragenomic FCGR analysis suggested by this study. Another significant observation of this study, based on deviations in the genomic signature observed in the genomes of eukaryotic species such as human and maize, is that not all randomly selected genomic segments accurately reflect the species-specific genomic signature. This observation is especially important for downstream applications such as phylogenetic inference and sequence classification, and it highlights the significance of the proposed computational pipelines for the selection of a DNA representative genomic segment that can serve as reliable genome proxy.

This study opens several avenues for future research. Since FCGRs are *k*-mer-based representations of DNA sequences, their effectiveness can vary depending on both sequence length and the selection of parameter *k*. Given that the optimal choice of *k* is highly dataset-dependent, with no single value consistently outperforming others across all datasets [84], exploring adaptive or data-driven strategies for parameter selection could further improve consistency of the FCGR based analysis [84]. Moreover, while FCGR-based methods offer a computationally efficient and alignment-free alternative to traditional approaches like BLAST [4], they produce numerical dissimilarity scores without direct biological interpretation [71]. Future efforts could aim to correlate these quantitative measures with interpretable inferences [71]. Additionally, the representative selection pipelines and intragenomic variation experiments presented in this study have been evaluated on four complete genomes but could be extended to the study of species with potentially different characteristics, including those with bloated genomes where repetitive sequences dominate (e.g., South American lungfish, *Lepidosiren paradoxa* [65]). Finally, while our experiments suggest that DSSIM is the most effective distance measure for comparing FCGRs, one could leverage machine learning to develop new distance measures that could potentially align more closely with biological factors.

Overall, this study provides visual and quantitative methods, as well as a software tool, for efficient intragenomic genome signature analysis, with potential applications in detecting and visualizing spatial variation, revealing genomic islands [58], detecting horizontal gene transfer events [7, 17], exploring structural and organizational features [40], and investigating evolutionary biology hypotheses [10] that cannot be tested through traditional homology-based analyses.

## Acknowledgements

The authors declare financial support was received for the research, authorship, and/or publication of this article. This work was supported by the Natural Science and Engineering Research Council of Canada Grants RGPIN-2022-03547 to G.S.R., RGPIN-2023-05256 to K.A.H. and RGPIN-2023-03663 to L.K. This research was enabled in part by support provided by Digital Research Alliance of Canada RPP (Research Platforms Portals), https://alliancecan.ca/, Grant 616 to K.A.H. The funders had no role in the preparation of the manuscript.

## Author contributions statement

N.S. and L.K. conceived the study and wrote the manuscript. N.S. assisted by the co-authors designed the experiments and N.S. and G.S.R. performed the experiments. All authors conducted the data analysis and edited the manuscript, with K.A.H. contributing biological expertise and C.d.S contributing statistical expertise. All authors read and approved the final manuscript.

## Availability of the datasets

The datasets generated and/or analyzed during the current study are all available in public repositories, and the links can be found in Fig. 1b or associated literature. The *CGR-Diff* tool developed for this study, along with all datasets used, is available at https://github.com/Niousha12/Intragenomic_analysis.

## A. Detailed Methods

### A.1 Distance Measures Analyzed

In the calculation of vector-based distances, ??_*i*_ represents the *i*-th position in the FCGR vector *X*, ??_*i*_ represents the *i*-th position in the FCGR vector *Y*, and *L* denotes the length of the vectors, which is equal to the total number of pixels in an FCGR image.

**Normalized Euclidean distance** is a vector-based distance and is the normalized derivation of the Euclidean distance that is more suitable for comparison across different scales. In our study, to compute the Normalized Euclidean distance between two FCGR vectors, we use the following equation:

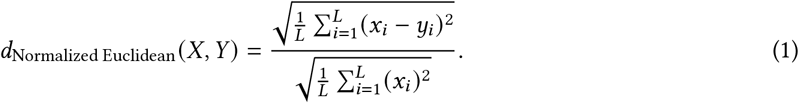

In this formulation, one of the FCGR images is used as the reference for normalization. A disadvantage of this distance when comparing two FCGRs is its lack of symmetry; specifically, *d*_Normalized Euclidean_ (*X, Y*) is not necessarily equal to *d*_Normalized Euclidean_ (*Y, X*). To address this issue, we compute the Euclidean norm (i.e., the denominator of Eq. 1) for both FCGRs and use the image with the larger value as the reference. A value of 0 in Normalized Euclidean distance indicates that the two images are identical, while a value in the range (0, 1] indicates that the two images are somewhat different, with the error being within the range of the reference image’s magnitude. However, a value greater than 1 indicates that the two images are significantly different, with the error exceeding the range of the reference image’s magnitude.

**Cosine distance**, which is another vector-based distance, is calculated by subtracting the cosine similarity of two FCGR vectors from 1. Considering that cosine similarity takes a value between [−1, 1], the cosine distance theoretically varies between [0, 2]. In order to calculate the cosine similarity and then cosine distance we can use the following equations:

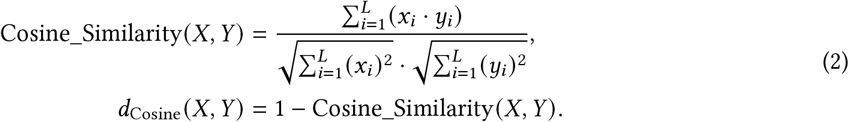

**Manhattan distance** is another vector-based distance measure. It is computed as the sum of the absolute differences between corresponding elements of two FCGR vectors, and it can theoretically range from 0 to∞. However, in our case, the upper bound is limited to 2^*k*^ × 2^*k*^ due to the resolution of the FCGRs, where *k* represents the value used in the *k*-mer. The Manhattan distance between two FCGR vectors is given by:

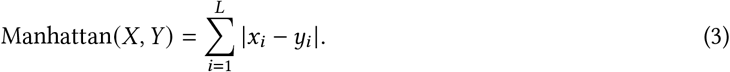

**Structural Dissimilarity Index (DSSIM)**, is calculated by subtracting Structural Similarity Index (SSIM) from 1. SSIM is a measure used to quantify the similarity between two images in three main aspects: luminance, contrast, and structure [77]. Luminance comparison measures the difference in the mean intensity or brightness of the images, contrast comparison measures the differences in the standard deviation or contrast of the images, and structure comparison measures the correlation between the local structures and patterns of the images. The SSIM between two images *X* and *Y* combines these three comparisons into a single score:

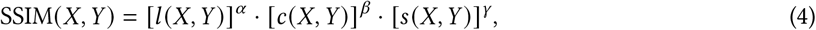

where *l* (*X, Y*) is the luminance comparison, *c* (*X, Y*) is the contrast comparison, and *s* (*X, Y*) is the structure comparison function between the two images. The parameters *α, β*, and *γ* are used to adjust the importance of each term.

Luminance measures the brightness level of an image. The luminance comparison function evaluates how the mean intensity (brightness) values of the two images differ. Given two images *X* and *Y*, the luminance comparison function is defined as:

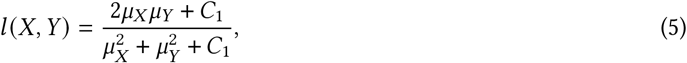

where *μ*_*X*_ is the local mean intensity matrix of image *X, μ*_*Y*_ is the local mean intensity matrix of image *Y*, and *C*_1_ is a small constant to avoid division by zero. Typically, *C*_1_ = (*K*_1_*L*) ^2^, where *L* is the dynamic range of the pixel values and *K*_1_ is a small constant, e.g., 0.01. The local mean of image *X* is computed as:

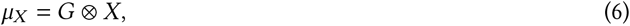

where *X* is the input image and *G* is a two-dimensional Gaussian kernel with a variance of 1.5 that is convolved with *X*. Thus, *μ*_*X*_ is a matrix of the same size as the input image, containing a blurred version of *X*.

The function *l* (*X, Y*) is designed to be 1 when the luminance of the two images is the same and decreases as the luminance difference increases.

Contrast measures the difference in the intensity values within an image, reflecting how varied the pixel values are from the mean. The contrast comparison function between two images *X* and *Y* is defined as:

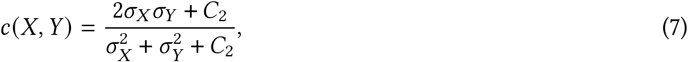

where *σ*_*X*_ and *σ*_*Y*_ are the local standard deviation of the corresponding images *X* and *Y*, which represent the image contrast. Additionally, *C*_2_ is another small constant to avoid division by zero. Typically, *C*_2_ = (*K*_2_*L*)^2^, where *K*_2_ is another small constant, e.g., 0.03, and *L* is the dynamic range of the pixel values.

To compute the local standard deviation of an image, we can convolve a Gaussian kernel filter using the following formulation:

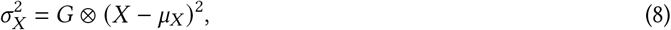

where *G* represents the Gaussian kernel filter and denotes convolution. This function is designed to be 1 when the contrast of the two images is the same and decreases as the contrast difference increases.

Structure measures the correlation between the patterns of pixel intensities in the images, reflecting how similar the local structures are. The structure comparison function between two images *X* and *Y* is defined as:

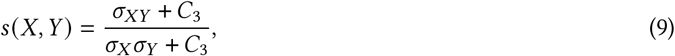

where *σ*_*XY*_ is the local covariance of images *X* and *Y*, representing how well the pixel intensities in one image can be predicted by the pixel intensities in the other. *C*_3_ is a small constant, typically *C*_3_ = *C*_2_/2.

The local covariance can be computed by convolving the product of the deviations from the mean of the two images with a Gaussian kernel filter:

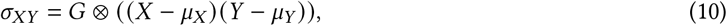

where *G* represents the Gaussian kernel filter and ? denotes the convolution operation. This function is designed to be 1 when the structural patterns of the two images are perfectly correlated and decreases as the correlation weakens.

The overall SSIM is computed by combining these three components as mentioned in Equation 4. To obtain the structural dissimilarity index (DSSIM), we subtract the SSIM from 1. SSIM values range from -1 to 1, where 1 indicates perfect similarity between the images, 0 indicates no similarity, and values less than 0 indicate structural dissimilarity. Consequently, the SSIM distance (*d*_SSIM_), defined as 1 − SSIM, ranges from 0 (for identical images) to 2 (for images that are completely dissimilar or negative of each other).

**Descriptor distance** between two images refers to a measure of similarity or dissimilarity based on specific features extracted from those images. After the features are extracted, they are typically converted into a numerical vector known as descriptors. The distance is then computed as the Normalized Euclidean distance between the corresponding descriptor vectors of the two images. In the case of FCGR comparison, to obtain the descriptor vector, we follow hierarchical image descriptors suggested by [33, 41].

For two FCGR images *X* and *Y*, where 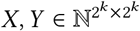, we calculate the distance as follows: first, we split each of the two images into non-overlapping sub-images of size 2^*m*^ × 2^*m*^, where *m* (*m < k*) is a hyperparameter of the distance measure [33]. After this division, we have 2^2(*k*−*m*)^ submatrices *X*_*i j*_ and *Y*_*i j*_ with *i, j* = {1, …, 2^*k*−*m*^} [33]. To create the descriptor vector for a sub-image *X*_*i j*_, we first divide the range of possible pixel values in an FCGR image into *r* bins. These bins are defined as [0, *k*_1_), [*k*_1_, *k*_2_), …, [*k*_*r* −1_, ∞). Next, we construct a vector of length *r*, (*b*_1_, *b*_2_, *b*_3_, …, *b*_*r*_), where each component *b*_*z*_ represents the total number of pixels within the sub-image *X*_*i j*_ whose values fall into the corresponding bin [*k*_*z*−1_, *k*_*z*_). Therefore, the extracted features from the images are local histograms with predefined bins. The final descriptor vector (DV) for the whole FCGR image is obtained by concatenating the descriptor vectors of all sub-images [33]. Finally, we use the following formulation for calculating the descriptor distance in our study:

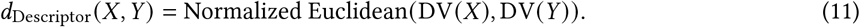

Similar to Normalized Euclidean, a descriptor distance of 0 means that the descriptor vectors of two images are completely identical, while values greater than 1 indicate a significant distance between the descriptor vectors.

**Learned Perceptual Image Patch Similarity (LPIPS)** is a deep learning-based method for measuring the similarity and distance between two images [83]. This method utilizes the intermediate feature space of a convolutional neural network to produce a distance measure that considers both structural and perceptual similarities of two images.

The input image passes through all these layers sequentially, with each layer extracting specific features. These layers are trained to recognize high-level features and textures in images. The early layers focus on structural features, while the later layers capture more perceptual aspects. To calculate the distance, LPIPS first computes the Euclidean distance between the feature maps of the two input images at corresponding layers. These layerwise distances are then weighted and summed to produce the final distance score.

To compute the LPIPS distance for two FCGR images, the same process is applied. Equation 12 shows the formulation for the LPIPS distance. In this equation, *X* and *Y* represent the two FCGR images, 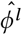 denotes the normalized feature map at layer *l* extracted using the pre-trained network, and *w*^*l*^ represents the learned weights used to combine the distances from *M* different layers.

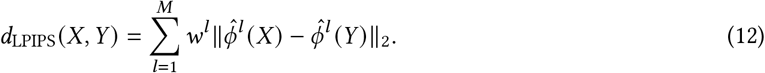

**Kolmogorov-Smirnov (K-S) distance** is based on the non-parametric test statistic that compares the distributions of two univariate datasets to determine if they differ significantly. As discussed earlier, one way to conceptualize FCGR representations is to view them as distributions of different *k*-mers. We can treat FCGRs as probability distributions of *k*-mers, allowing us to apply statistical methods to compare two FCGRs. This approach leverages the principles of probability and statistics to analyze and measure the similarities and differences between the FCGR representations. Therefore, we consider the K-S test statistic as a measure of distance between two FCGR images.

For two FCGR images 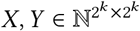, we initially convert each of them to one-dimensional vectors. Each vector is then converted to probability values by dividing each position value by the sum of all values. Next, we calculate the cumulative distribution function (CDF) of the probabilistic FCGRs. Finally, we return the maximum value over all possible values in the absolute subtraction of the CDF of the two images. The equation is:

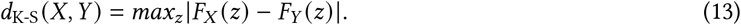

In this formulation, *F*_*X*_ (*z*) is the cumulative distribution function (CDF) of the probabilistic FCGRs at position *z*. The range of the K-S distance is [0, 1]. A value of 0 indicates that the empirical distribution functions of the two samples are identical, while a value of 1 indicates the maximum possible difference between the empirical distribution functions.

**Wasserstein distance** is a non-parametric method for comparing the distributions of two datasets to determine their similarity, analogous to the K-S test. More specifically, the Wasserstein distance measures the minimum amount of “work” required to transform one distribution into another. In this context, “work” is defined as the amount of distribution mass that needs to be moved multiplied by the distance it needs to be moved.

To apply the Wasserstein distance between two FCGR images *X* and *Y*, similar to the K-S test, we first flatten each FCGR image into a vector, then calculate the probability values, and subsequently utilize the cumulative distribution function (CDF). Finally, we use the following equation to compute the distance:

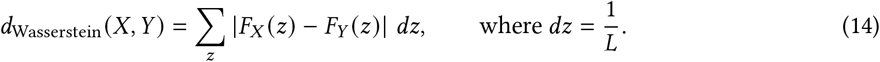

In this formulation, *F*_*X*_ (*z*) represents the cumulative distribution function (CDF) of the probabilistic FCGR *X* at position *z*, and *L* is the total number of positions or pixels.

The Wasserstein distance ranges from 0 to 1. A distance of 0 indicates that the two distributions are identical, meaning no “work” is needed to transform one distribution into the other. Conversely, a distance of 1 indicates that the two distributions are completely different, meaning they are as far apart as possible at every point *z*.

Before calculating the values of different distance measures on the FCGR images, we apply a preprocessing step to each of our distance groups. For image-based and vector-based distance measures, we apply the min-max normalization to rescale the FCGR values to the [0, 1] range, thereby enhancing the comparability of two FCGR images. The effectiveness of this normalization is well-documented in the literature, as it manages the large variance in FCGR images [45] and ensures that all elements contribute equally to the distance calculations [21]. A key advantage of this normalization in FCGR comparison is that it enables the comparison of FCGRs derived from sequences of different lengths. The normalization process for these two distance groups is applied as follows:

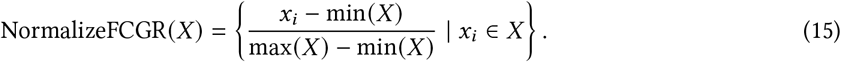

For probability-based distances, such as the K-S test and Wasserstein distances, which use statistical methods operating on probability distributions, we apply probability normalization to the FCGRs. This ensures that the sum of all values equals one before performing these tests. The normalization process is applied as follows:

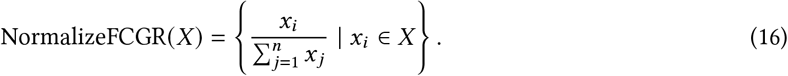

### A.2 Experiments for Human Intragenomic Distance Analysis (Exp 2.1)

The detailed description of the experiments conducted for the *Human Intragenomic Distance Analysis* are as follows:

- *Telomere vs. Telomere*: This experiment compares the distance between telomeric regions of different chromosomes. Telomeres are identical short sequence tandem repeats located at the ends of the p-arms and q-arms of all of the chromosomes [67]. The composition and structure of telomeres are known to be similar across different human chromosomes, and these similarities are attributed to the conserved nature of telomeric DNA sequences and the associated proteins that form these protective structures [44]. Human telomeres typically consist of thousands of short ‘TTAGGG’ tandem repeats, which vary in length with age, leading to telomere lengths that typically range from 5 to 15 Kbp [79]. These repeats are crucial for maintaining chromosome stability and integrity [44]. In this experiment, we calculate the average distance between the p-arm telomere of the first human chromosome and the p-arm telomeres of the other chromosomes. This approach can be applied to both p-arm and q-arm telomeres; however, for consistency, we choose to focus on the p-arm telomere. Similarly, this experiment could be extended to centromeres, as they also contain highly repetitive sequences of identical repeats [23], known as alpha-satellite DNA [70]. These alpha-satellite monomers are approximately 171 bp in length [70]. However, given the expected similarities in centromere comparisons, we focus exclusively on telomeres.
- *Heterochromatin vs. Heterochromatin*: Heterochromatin in all chromosomes consists of highly condensed, repetitive, and transcriptionally inactive regions that show similarities across different chromosomes [60, 72]. These regions are represented by black and three shades of gray in the NCBI Data Viewer, where darker shades correspond to more condensed chromatin, reflecting higher levels of chromatin compaction and staining intensity [22]. In this experiment, we initially extract the most condensed regions of different chromosomes, which are colored in black. For chromosomes 16 and 21, where only one black region is present, we also include the dark gray region. For chromosomes 15, 17, 19, 20, 22, and Y, where there are no black regions, we select the next most condensed region, represented by other shades of gray. For each chromosome, we calculate the average distance between the most condensed heterochromatic region distal to the centromere on the p-arm (or, if none exists, proximal to the centromere on the q-arm) and other heterochromatic regions within the same chromosome. We then report the average distance across all chromosomes and evaluate the performance of different distance measures.
- *Heterochromatin vs. Euchromatin*: Euchromatin describes regions of DNA that are less condensed and contain genes that are actively transcribed into RNA [8]. Euchromatin and heterochromatin regions differ significantly in their overall structure and function [8]. Euchromatin is associated with gene-rich regions and active transcription, while heterochromatin is linked to gene-poor regions and gene silencing [8]. Euchromatin has higher C+G content and CpG density, associated with active gene transcription, while heterochromatin has lower C+G content and fewer, often methylated, CpG sites, leading to gene silencing [52]. In this experiment, for each chromosome, we first measure the distance between the most condensed heterochromatic region distal to the centromere on the p-arm (or, if none exists, proximal to the centromere on the q-arm) and each of four randomly selected euchromatic segments. We then report the average distances across all chromosomes for comparison between the distance measures.
- *p-arm vs. q-arm (for acrocentric chromosomes)*: Acrocentric chromosomes are distinguished by their centromere being positioned very close to one end, resulting in a short p-arm and a long q-arm [69]. The short arm of these chromosomes is a stretch of DNA sequence that contains tandem repeat sequences, while their long arm contains less of these tandem repeat sequence arrays [49]. Large tandem repeats in chromosomes are DNA sequences composed of multiple copies of a particular sequence arranged in a head-to-tail (tandem) manner [20]. Inspired by the structural differences between the p-arm and q-arm of acrocentric chromosomes, we design this experiment using the acrocentric chromosomes in humans, which are the five autosomal chromosomes 13, 14, 15, 21, and 22 [49], as well as chromosome Y [16]. The repetitive region on the Y chromosome is located on the long arm (q-arm) and is fundamentally different in overall length and the length and nature of the tandem repeat arrays [16]. In our experiment, we examine each of these chromosomes individually. For each chromosome, we randomly select a 500 Kbp segment from the p-arm and another 500 Kbp segment from the q-arm, then calculate the distance between them. This process is repeated 100 times to ensure variability and randomness in segment selection, and the results are then averaged.
- *Large Tandem Repeat Arrays*: Large tandem repeats, classified as moderately repetitive structures [42], vary significantly in size and the number of repeat units and are categorized based on their length and organization [78]. These repeats can be found in the p-arm of acrocentric chromosomes and the q-arm of chromosome Y. However, the large tandem repeat arrays in the q-arm of chromosome Y differ in both length and structural composition from those in acrocentric chromosomes [64]. In this intragenomic experiment, we compare the q-arm of the Y chromosome to the large tandem repeat arrays of the acrocentric chromosomes by calculating the distance between the cytoband q12 of chromosome Y and each of the cytobands containing large tandem repeat arrays on chromosomes 13, 14, 15, 21, and 22. The average of these distances is then reported. The q12 cytoband on the Y chromosome is approximately 35 Mbp in length, which is part of the total 62 Mbp of the Y chromosome. The approximate lengths of the large tandem repeat arrays are as follows: for chromosome 13, 16 Mbp out of 114 Mbp; for chromosome 14, 10 Mbp out of 101 Mbp; for chromosome 15, 17 Mbp out of 100 Mbp; for chromosome 21, 11 Mbp out of 45 Mbp; and for chromosome 22, 13 Mbp out of 51 Mbp. Compared to the *p-arm vs. q-arm* experiment, which contrasts tandem repeats with non-repeats, the *Large Tandem Repeat Arrays* experiment focuses on comparing different types of tandem repeat regions.
- *Arbitrary Sequences*: In the final experiment, we aim to determine an intermediate intragenomic distance between FCGRs. To do so, we randomly select two non-overlapping 500 Kbp sequences from a randomly chosen chromosome and compute the distance between their FCGRs. This process is repeated 100 times, and the average distance is used for comparison. Given the 100 repetitions of this random selection process, we expect it to sample diverse combinations of genomic sequences, including tandem repeat arrays and non-repetitive regions.

## B Supplementary Figures

**Figure S1:**
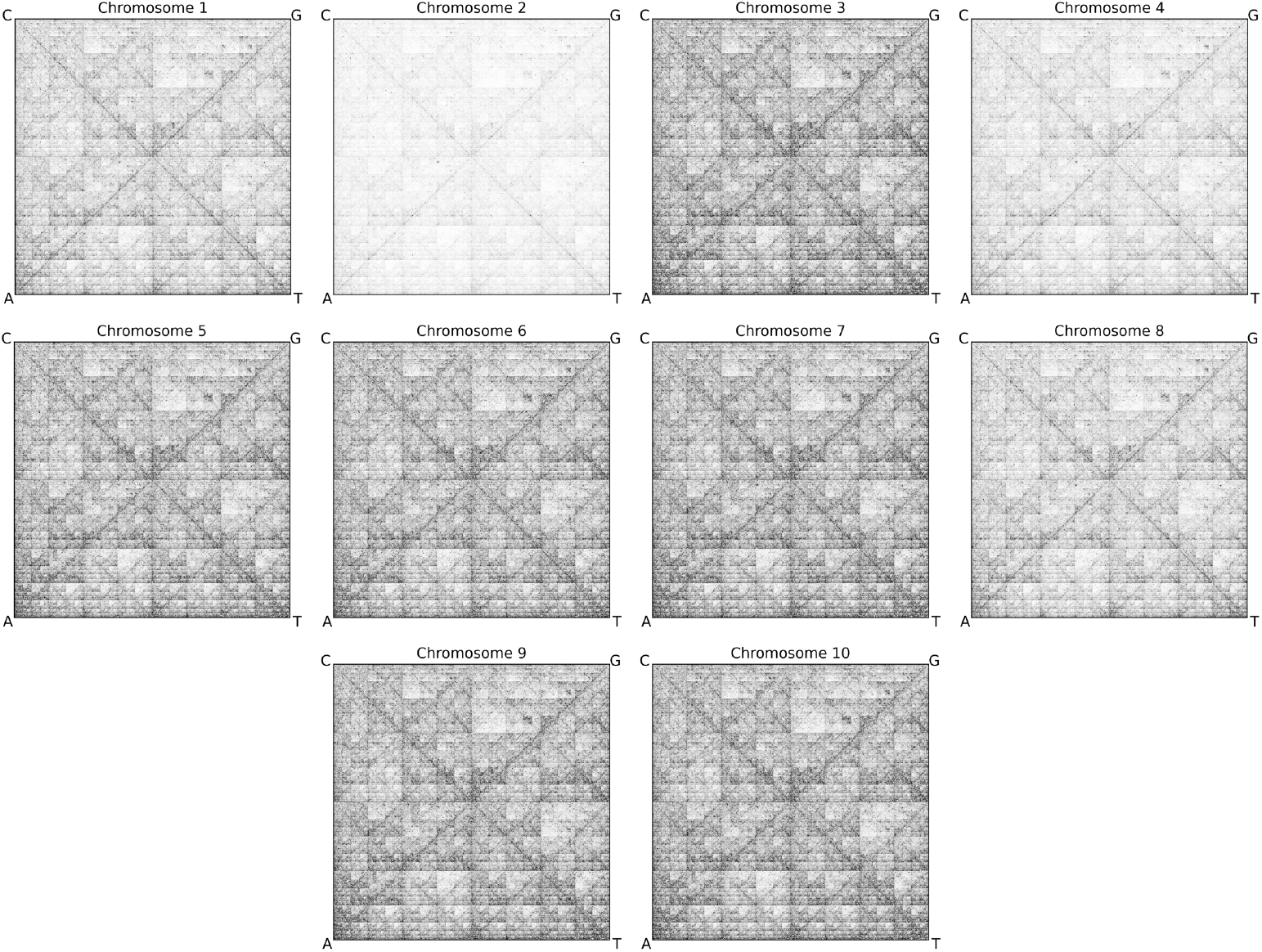
Analyses of the Maize Genome Genetic Signature. Each image displays the FCGR of a complete maize chromosome, generated using *k* = 9. The patterns reveal both conserved genomic features and variations in *k*-mer distribution, indicating that while the overall structure is largely consistent across chromosomes, differences in shading highlight regions with uneven *k*-mer composition. Notably, chromosomes 2 and 4 appear paler, suggesting the presence of highly repeated *k*-mers (9-mers in this case) within their sequences.

**Figure S2:**
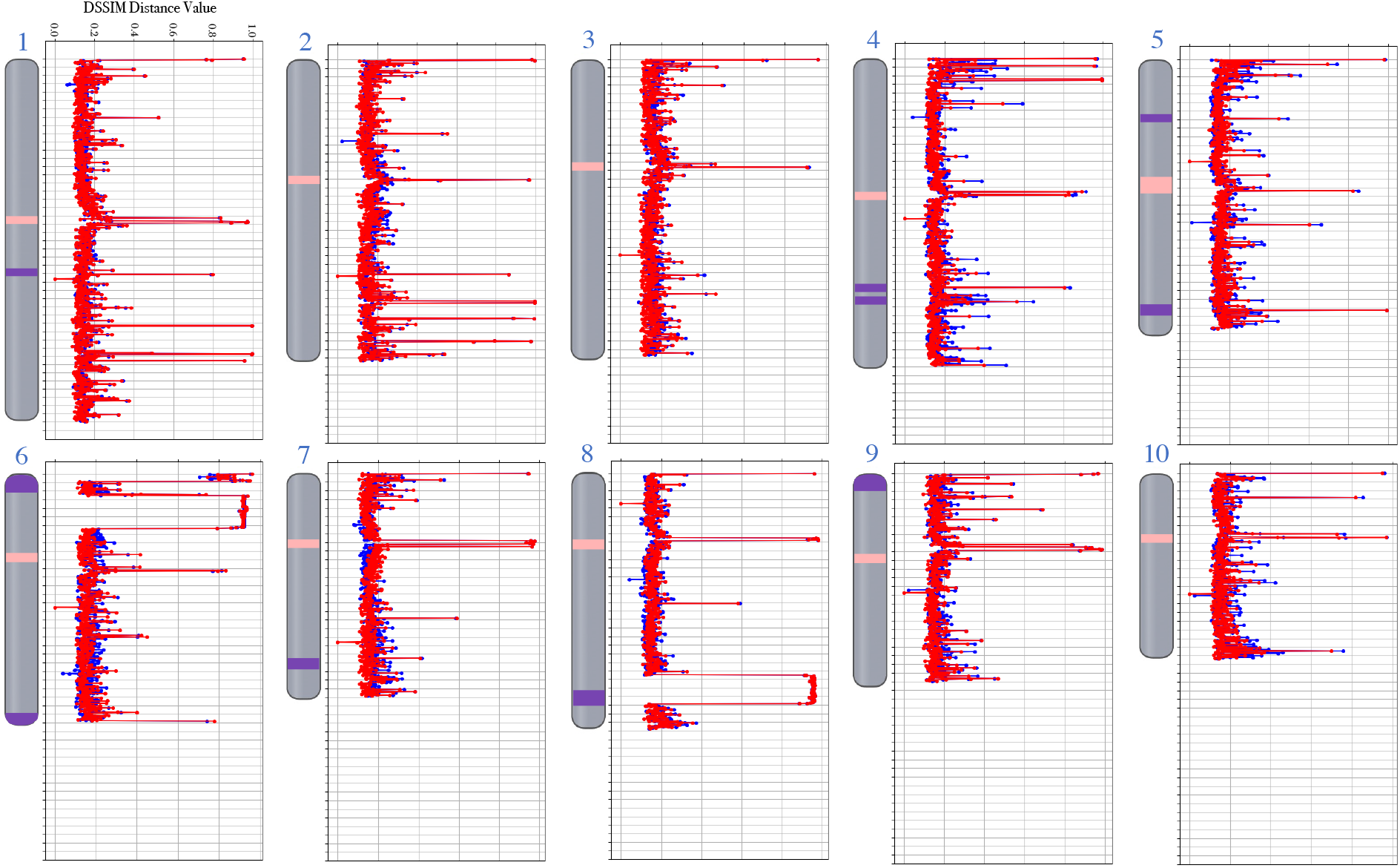
DSSIM distances of consecutive segments from the representative segment across all maize chromosomes. The red and blue line plots represent Exp 3.1 and Exp 3.2, respectively. Each point corresponds to the calculated distance between a 500 Kbp genomic segment and the representative segment selected by the proposed pipelines. The horizontal axis displays DSSIM distance values ranging from 0 to 1 in increments of 0.2, where higher values indicate greater dissimilarity. The vertical axis is segmented into intervals of 20, corresponding to sequential 500 Kbp segments along each chromosome. To the left of each plot, chromosome ideograms are generated using the NCBI Genome Data Viewer [55], and the suggested centromere and KNOB180 region annotations by Hufford et al. [26]. These ideograms provide an approximation of the positions of centromeres (pink regions) and tandem repeat arrays known as KNOB180 (purple regions).

**Figure S3:**
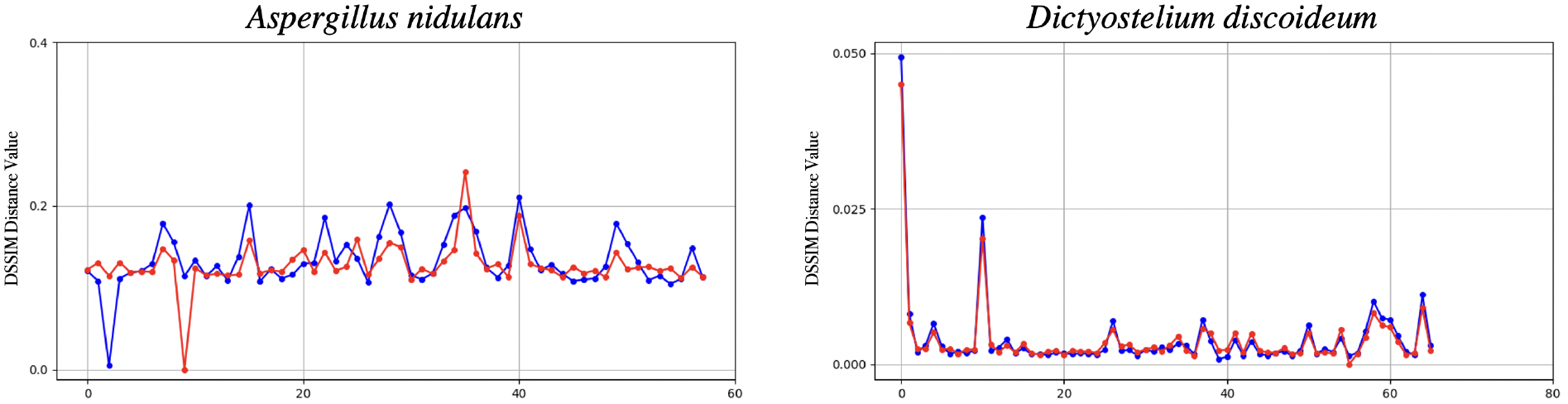
DSSIM distances of consecutive segments from the representative segment for two eukaryote species (left: *Aspergillus nidulans*, right: *Dictyostelium discoideum*). The red line plots show the DSSIM distances between consecutive 500 Kbp segments and the representative segment identified by RepSeg, while the blue line plots illustrate the DSSIM distances from the representative segment selected by aRepSeg. The horizontal axis is segmented into intervals of 20, representing sequential 500 Kbp segments along each chromosome, while the vertical axis shows the DSSIM distance values. Notably, the vertical axis range differs between the two species to accommodate their respective DSSIM distance distributions. In *Aspergillus nidulans*, DSSIM distances are higher, reaching up to 0.4, whereas in *Dictyostelium discoideum*, distances remain much lower, with a maximum of 0.05.

**Figure S4:**
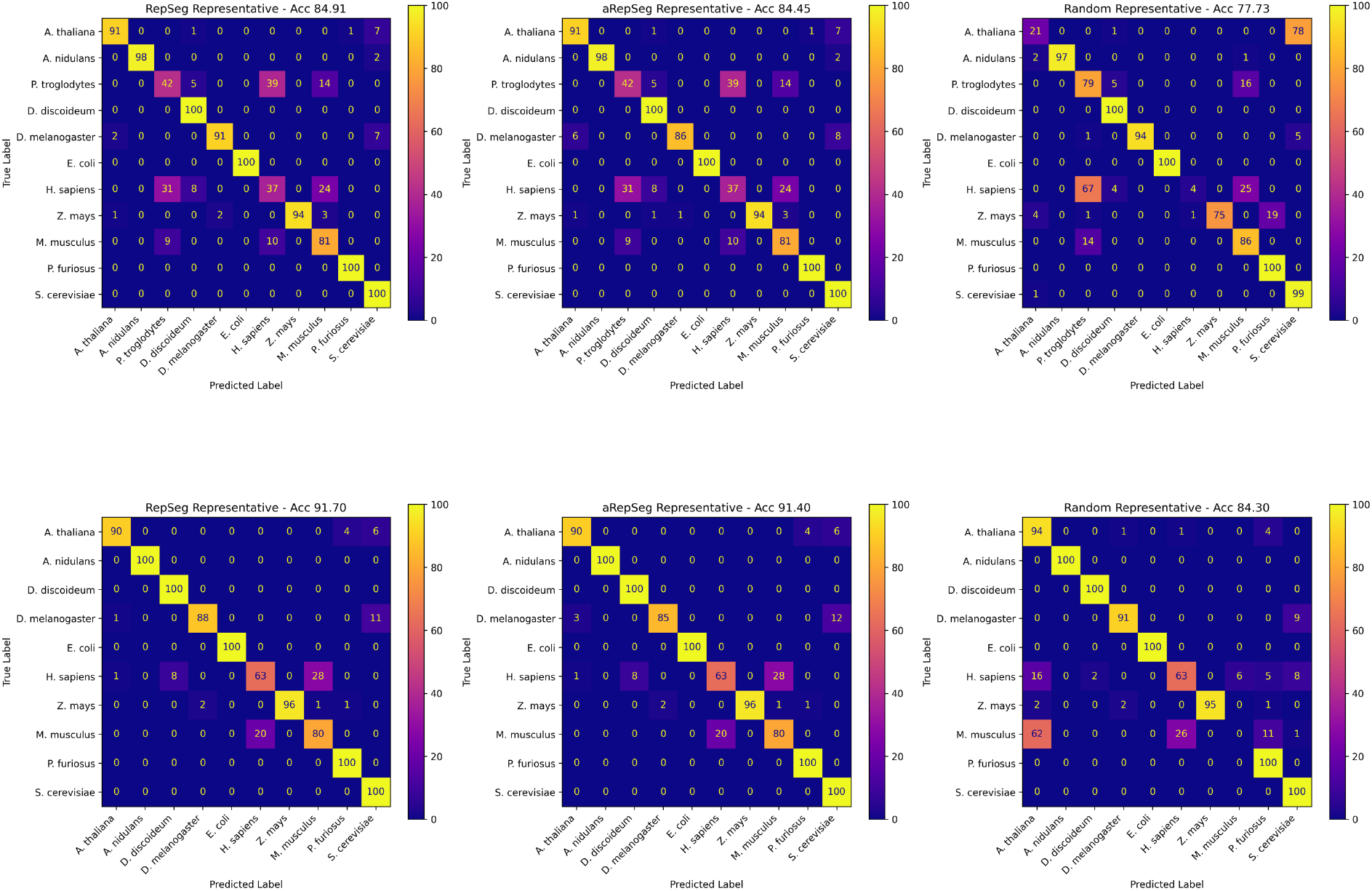
Confusion matrices for the nearest neighbor classifier (Experiment 4). The top row displays confusion matrices for the experimental setup that includes all species from the study. Due to the high genetic similarity between the human genome and chimpanzee genome, most misclassifications occur between these two species. The bottom row illustrates the results when the chimpanzee species is excluded from the test, leading to improved classification performance. Notably, while the overall accuracy increases in this scenario, the difference in accuracy between the pipeline-selected representative and the random representative remains the same. Among the approaches, the RepSeg method (left) achieves the highest accuracy, with aRepSeg (middle) performing similarly well, whereas the random representative (right) is less optimal in comparison.

## C Supplementary Tables

* Distance measures: (1) Normalized Euclidean distance, (2) Cosine distance, (3) Manhattan distance, (4) Structural Dissimilarity Index (DSSIM), (5) Descriptor distance, (6) Kolmogorov-Smirnov (K-S) distance, (7) Wasserstein distance, and (8) Learned Perceptual Image Patch Similarity (LPIPS).

*Among all the species in this table, the assembly for *Paramecium caudatum* is at the scaffold level.

**Table S1:**
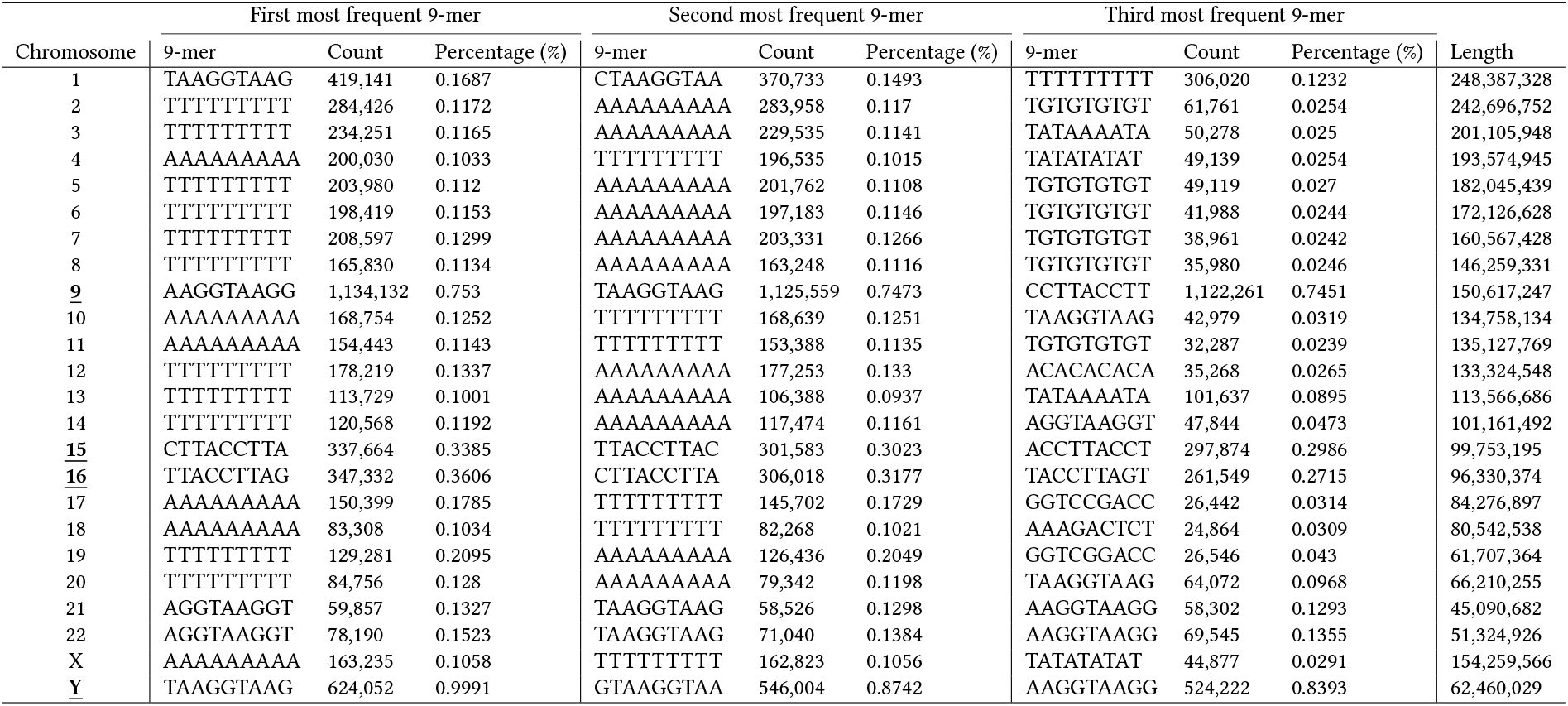
Most frequent 9-mers across all human chromosomes, including their counts and percentages. The table reports the top three most frequent 9-mers identified in each human chromosome (1–22, X, and Y), based on their raw occurrence counts and the percentage each represents out of the total number of 9-mers in that chromosome. These values are shown alongside the total length of each chromosome. Notably, 9-mers such as TTTTTTTTT and AAAAAAAAA appear frequently across multiple chromosomes, while chromosomes 9 and Y exhibit exceptionally high occurrences of specific 9-mers, such as AAGGTAAGG and TAAGGTAAG, respectively. The <u>**bolded**</u> chromosomes indicate those with lighter FCGR images.

**Table S2:**
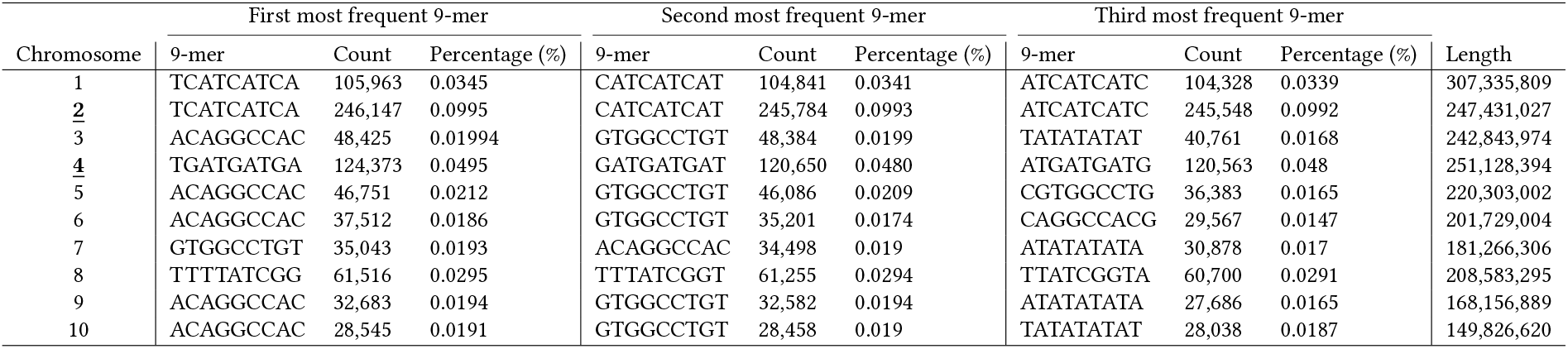
Most frequent 9-mers across all maize chromosomes, including their counts and percentages. The table lists the top three most frequent 9-mers found in each maize chromosome (1–10), reporting both their raw counts and the percentage they consist of the total number of 9-mers within the respective chromosome. These values are shown alongside the total length of each chromosome. Notably, chromosomes 2 and 4 display particularly high occurrences of specific 9-mers. The <u>**bolded**</u> chromosomes indicate those with lighter FCGR images.

**Table S3:** The *p*-values from the Wilcoxon signed-rank test for various distance measures, comparing the human-human distances with the distances between human and each species in the corresponding row. It is expected that no significant difference (i.e., a *p*-value greater than 0.05) will be observed when comparing human-human distances with human-*P. troglodytes* and human-*M. musculus*, as they are both from the Mammalian class. However, the Manhattan, Descriptor, and Wasserstein distance measures yield values smaller than 0.05 when comparing human-human distances with human-*M. musculus*, which does not support the expectation of no significant difference among mammals.

| human-human vs.<br>human-other species | Normalized<br>Euclidean | Cosine | Manhattan | Descriptor | DSSIM | LPIPS | K-S | Wasserstein |
| --- | --- | --- | --- | --- | --- | --- | --- | --- |
| *P. troglodytes* | 0.1507 | 0.4745 | 0.1648 | 0.2355 | 0.1777 | 0.3919 | 0.3348 | 0.9124 |
| *M. musculus* | 0.1161 | 0.6352 | 9.0e-4 | 0.0348 | 0.0961 | 0.9206 | 0.6799 | 0.0415 |
| *D. melanogaster* | 6.75e-10 | 4.67e-11 | 1.79e-11 | 6.99e-7 | 1.11e-14 | 6.00e-8 | 8.94e-16 | 0.0336 |
| *S. cerevisiae* | 1.10e-6 | 2.81e-8 | 5.01e-11 | 0.0256 | 3.54e-13 | 5.89e-8 | 9.27e-14 | 0.0333 |
| *A. thaliana* | 3.69e-6 | 2.17e-8 | 2.04e-7 | 0.0059 | 1.09e-11 | 3.40e-6 | 1.24e-9 | 0.0684 |
| *P. caudatum* | 3.44e-15 | 3.24e-14 | 5.19e-15 | 9.88e-4 | 1.53e-14 | 2.26e-16 | 1.38e-15 | 2.26e-16 |
| *P. furiosus* | 1.75e-16 | 3.42e-12 | 3.90e-18 | 9.87e-16 | 1.15e-17 | 6.54e-17 | 1.73e-8 | 2.07e-4 |
| *E. coli* | 1.74e-17 | 1.27e-15 | 3.90e-18 | 1.91e-16 | 4.27e-18 | 4.17e-15 | 1.16e-14 | 2.73e-6 |

**Table S4:**
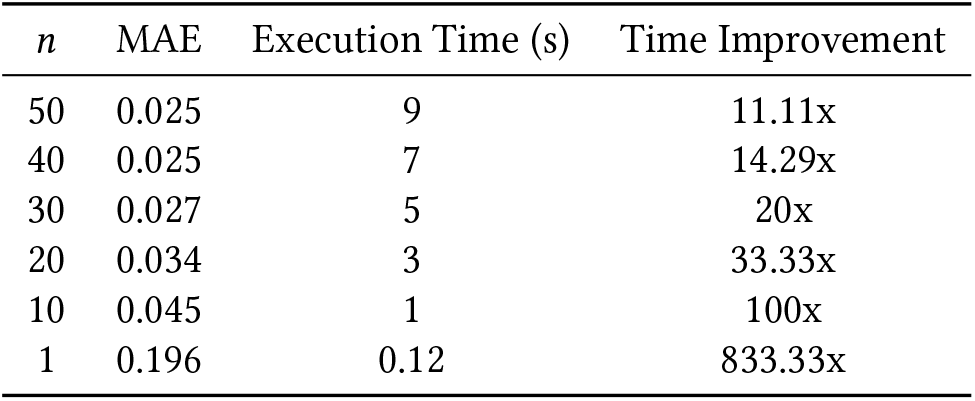
Effect of the hyperparameter *n* (the size of the set *Ŝ* in ARSSP) on runtime and mean absolute error (MAE) between ARSSP and RSSP for human chromosome 1. The MAE and execution time, averaged over 100 runs of the ARSSP pipeline, illustrate how changes in *n* influence both accuracy and computational cost. The time improvement, representing the relative speedup compared to the RSSP execution time (∼100s), highlights the trade-off between computational efficiency and accuracy, which provides insights into the performance of ARSSP in comparison to RSSP.

| $n$ | MAE | Execution Time (s) | Time Improvement |
| --- | --- | --- | --- |
| 50 | 0.025 | 9 | 11.11x |
| 40 | 0.025 | 7 | 14.29x |
| 30 | 0.027 | 5 | 20x |
| 20 | 0.034 | 3 | 33.33x |
| 10 | 0.045 | 1 | 100x |
| 1 | 0.196 | 0.12 | 833.33x |

